# Central Amygdala Neuronal Ensembles Coordinate Visceral Pain and Its Affective Behaviors

**DOI:** 10.1101/2025.07.21.666026

**Authors:** Vijay K Samineni, Julian N Sackey, Jun-Nan Li, Lite Yang, Alexander Chamessian, Sienna B Sewell, Hannah Hahm, Yufen Zhang, Bo Zhang, Robert W. Gereau

**Author notes:** Contact information: Robert W. Gereau IV, Vijay K Samineni.

## Abstract

Visceral pain and its affective component associated with cystitis remain poorly understood. Here, we delineate the role of central amygdala (CeA) neuronal ensembles in encoding and modulating cystitis-induced bladder pain and its affective components. Utilizing a multidisciplinary approach combining behavioral assays, optogenetic manipulations, whole-cell electrophysiology, and activity-dependent genetic labeling, we identified functionally discrete CeA subpopulations that are selectively recruited during bladder inflammation. Bidirectional optogenetic modulation of these ensembles produced opposing effects on nocifensive and anxiety-like behaviors, indicating their causal involvement. Single-nucleus RNA sequencing of FosTRAP-labeled neurons revealed distinct transcriptional signatures associated with inflammatory activation. Integrating electrophysiological and transcriptomic data, we demonstrate that cystitis drives cell type–specific adaptations in CeA circuits. These findings provide mechanistic insight and uncover a molecularly and functionally defined CeA ensemble that orchestrates the sensory and affective dimensions of visceral pain.

## Introduction

Chronic visceral pain syndromes such as interstitial cystitis/bladder pain syndrome (IC/BPS) are chronic pelvic pain conditions that disproportionately affect women worldwide (Berry et al 2011, Hanno et al 2015, Rosenberg et al 2007). Defined by persistent bladder-associated pain exacerbated by urinary filling, IC/BPS is frequently accompanied by urgency, frequency, and referred pain which cause profound disruption in quality of life. ^1–5^. However, the clinical burden of IC/BPS extends beyond somatic symptoms; a substantial proportion of patients experience comorbid affective disturbances, particularly anxiety, which potentiate the suffering and underscore a complex interplay between bodily pain and emotional dysregulation^6–13^. Traditionally conceptualized as peripheral urological dysfunction, IC/BPS is increasingly recognized as a disorder with extensive central nervous system involvement ^14–17^. The high prevalence of comorbid anxiety in IC/BPS patients implicates long lasting maladaptive plasticity in pain- and emotion-related brain circuits ^18–21^. Functional neuroimaging studies in IC/BPS patients reveal heightened activity in limbic regions such as the amygdala ^22–27^, suggesting that central amplification of pain signals may underlie both persistent visceral pain and associated psychological comorbidities.

The central nucleus of the amygdala (CeA) is well established sub-nuclei in the amygdala known for encoding sensory and emotional memories ^28–30^. Decades of work with behavioral, and physiological studies have consistently implicated the CeA as a key node of chronic pain processing ^31–45^. Functional imaging studies in patients with IC/BPS consistently reveal enhanced amygdala activity ^24,46,47^, while lesion and circuit-level manipulation studies in rodents have demonstrated that the CeA is necessary for the full expression of bladder hypersensitivity and its affective correlates ^36,37,48–50^. Notably, induction of cystitis by cyclophosphamide (CYP) triggers immediate early gene expression in the CeA ^51^, and optogenetic activation of this region recapitulates pain-like responses, suggesting that CeA output is not merely epiphenomenal but causally implicated in bladder pain. Our prior work demonstrated that optogenetic activation of the right CeA sensitizes bladder responses, implicating enhanced CeA output in the facilitation of bladder pain ^36^. Furthermore, recent studies have highlighted the influence of neuromodulatory peptides such as PACAP and CGRP, released from upstream regions like the parabrachial nucleus, on CeA activity in bladder pain states ^48,49^. Beyond its role in pain modulation, the CeA is also implicated in the regulation of fear, anxiety, and emotional memory ^52–61^. The intersection of these domains suggests that the CeA may serve as a neural convergence wherein sensory and affective components of visceral pain are encoded within overlapping neuronal ensembles. Yet despite this converging evidence, the functional and genetic identity of CeA neurons that link bladder pain to emotional suffering remains largely undefined.

Despite the well-established involvement of the CeA in both BPS and emotional processing, critical questions remain regarding its circuit-level contributions to IC/BPS. Specifically, it is unknown whether distinct CeA neuronal subtypes encode or modulate the sensory and affective components of cystitis-associated pain, whether cystitis induces long-lasting adaptive changes in CeA circuitry, and how these changes might contribute to the maintenance of bladder pain. Moreover, the molecular identity of CeA circuits that participate in cystitis-induced visceral hypersensitivity remains unclear. To address these questions, we deployed a mouse model of bladder pain induced by intraperitoneal injection of cyclophosphamide (CYP). This model produces the cardinal signs of IC/BPS including pain on bladder filling, referred hypersensitivity to the abdomen, and increased urinary frequency^62^. We used electrophysiology, activity dependent labelling, optogenetics and single cell RNA sequencing to systematically identify and characterize a genetically defined population of CeA neurons that modulates cystitis-induced bladder pain and associated negative affective states.

## Results

To examine whether cystitis modulates the intrinsic excitability of central amygdala (CeA) neurons, we performed whole-cell current-clamp recordings on neurons within the CeA in mice that received either intraperitoneal (i.p.) saline or CYP injections (Fig. 1A), with recordings conducted 4hrs post injection to match peak abdominal referred pain sensitization (Fig. S1). Consistent with previous reports, the majority of CeA neurons exhibited a ‘late-firing’ phenotype—95.8% (23/24) in saline-treated and 92.3% (24/26) in CYP-treated animals. However, we identified two intrinsically distinct subpopulations within the CeA, differentiated by their firing responses to incremental current injections (Fig. 1B, Fig. S2A). One subgroup displayed reduced firing at higher current amplitudes, which we termed ‘adapting neurons’ (Fig. 1B, left), while the second subgroup exhibited a continuous increase in action potential (AP) firing with increasing current, designated as ‘non-adapting neurons’ (Fig. 1B, right). Adapting neurons exhibited a significantly lower current threshold (Fig. S2B) and reached their peak firing frequency at lower current (APmax current; Fig. S2C) compared to non-adapting neurons in both saline- and CYP-treated mice, suggesting that adapting neurons either possess inherently higher excitability or or receive stronger excitatory or weaker inhibitory input. Moreover, adapting neurons demonstrated greater excitability than non-adapting neurons at lower current injections but lower excitability at higher current levels (Fig. S2A), indicating potential differences in their functional roles or activation thresholds under physiological conditions. In a variety of other intrinsic firing properties, little difference was observed between adapting and non-adapting neurons, though some properties were altered in neurons from CYP-treated mice (Fig. S2D-M). Despite these physiological differences, the relative proportions of adapting and non-adapting neurons remained consistent between saline- and CYP-injected groups (Fig. 1C). Collectively, these findings suggest that adapting and non-adapting CeA neurons constitute two physiologically distinct neuronal populations, potentially contributing differentially to the neural circuitry underlying bladder pain.

**Figure. 1.**
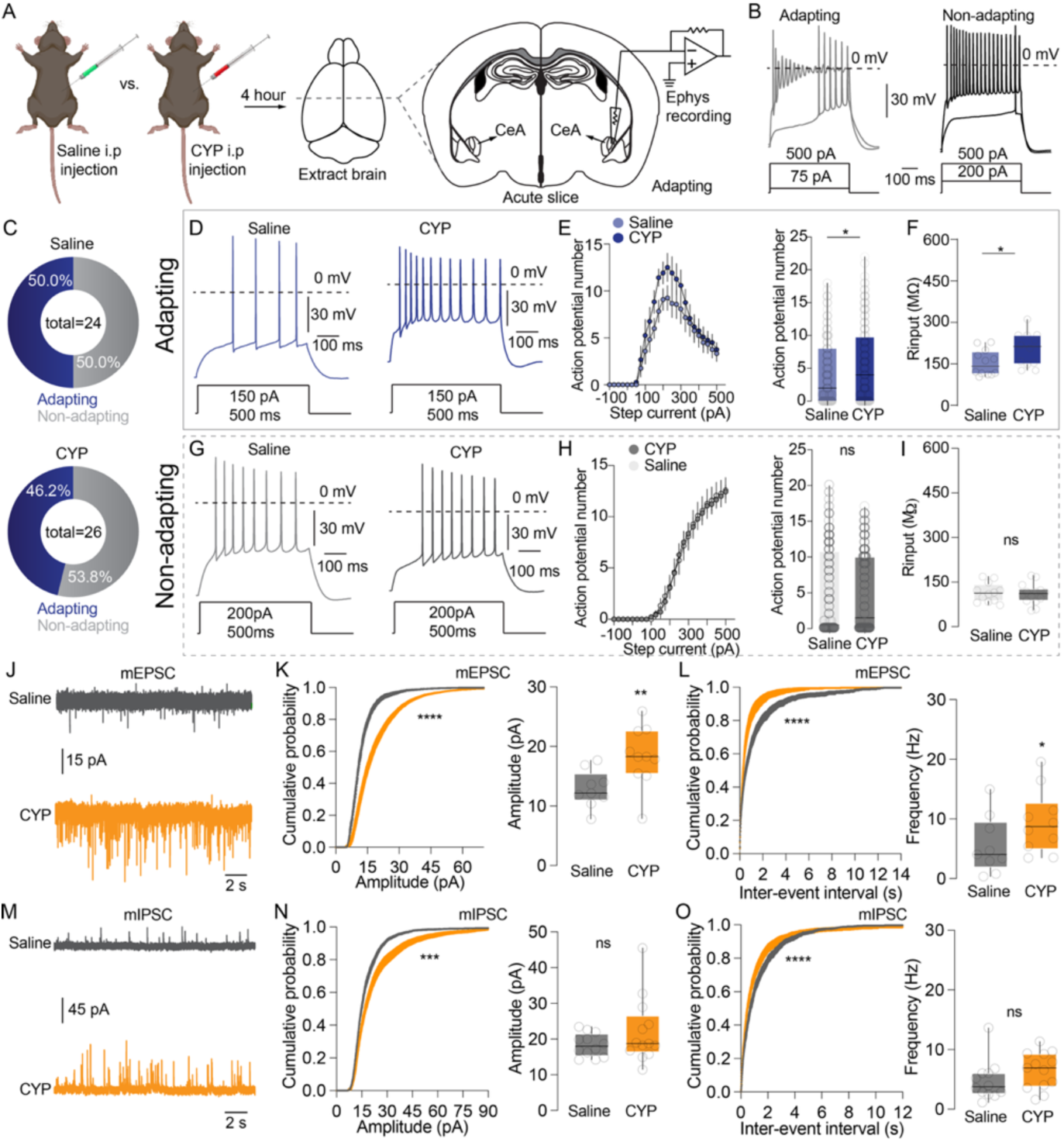
The alteration of neuronal excitability and synaptic plasticity of CeA neurons induced by cystitis. (A) Schematic illustrating the establishment of bladder pain model and the slice electrophysiology. (B) Current-clamp recording showing two intrinsically distinct neural types in CeA, adapting neuron showing decrease or cease of AP firing at higher current injection with 500ms duration and non-adapting neurons showing normal firing without decrease or cease of AP number at higher current injection. (C) The proportion of adapting and non-adapting CeA neurons in Saline-injected (top) and CYP-injected (bottom) mice. (D) Representative AP traces for adapting CeA neurons recorded from saline-injected (left) and CYP-injected (right) mice. (E) Plot of AP number vs. current injection for adapting CeA neurons from saline-injected and CYP-injected mice (left). Column bar graph displaying comparison of combined AP number (from - 100 pA to 500 pA) for adapting CeA neurons recorded from saline-injected and CYP-injected mice (right). Two-tailed Mann-Whitney test. (F) boxplot displaying comparison of input resistance for adapting CeA neurons recorded from saline-injected and CYP-injected mice. Two-tailed unpaired t test. (G) Representative AP traces for non-adapting CeA neurons recorded from saline-injected (left) and CYP-injected (right) mice. (H) Plot of AP number vs. current injection for non-adapting CeA neurons from saline-injected and CYP-injected mice (left). Column bar graph displaying comparison of combined AP number (from - 100 pA to 500 pA) for non-adapting CeA neurons recorded from saline-injected and CYP-injected mice (right). Two-tailed Mann-Whitney test. (I) boxplot displaying comparison of input resistance for non-adapting CeA neurons recorded from saline-injected and CYP-injected mice. Two-tailed unpaired t test. (J) Representative mEPSCs recorded from saline-injected (top) and CYP-injected (bottom) mice alongside quantifications of mEPSC amplitude (K) and frequency (L). (M) Representative mIPSCs recorded from saline-injected (top) and CYP-injected (bottom) mice alongside quantifications of mIPSC amplitude (O) and frequency (P). Kolmogorov-Smirnov test for (K, left), (L, left), (O, left) and (P, left), ***p < 0.001, ****p < 0.0001. Two-tailed unpaired t test for (K, right), (L, right) and (P, right). Two-tailed Mann-Whitney test for (0, right).

Next, we investigated whether bladder pain selectively alters the excitability of the two physiologically distinct CeA neuronal subpopulations. Our data revealed that CYP treatment significantly enhanced the excitability of adapting neurons, as evidenced by an upward shift in the input-output (current–AP number) relationship (Fig. 1E, left) and a significant increase in the total number of action potentials generated across a range of current injections (–100 pA to +500 pA) compared to saline-treated controls (Fig. 1D, 1E, right). This enhancement was accompanied by a significant increase in input resistance (R_input) specifically in adapting neurons following CYP administration. In contrast, the intrinsic membrane properties of non-adapting CeA neurons remained unchanged in response to CYP treatment (Fig. 1G– I), indicating a cell-type-specific effect of cystitis-induced pain on neuronal excitability. To further assess whether these changes in excitability were driven by altered synaptic inputs—and to distinguish between pre- and post-synaptic mechanisms—we performed whole-cell voltage-clamp recordings to measure miniature excitatory and inhibitory postsynaptic currents (mEPSCs and mIPSCs, respectively) in CeA neurons. CYP-induced cystitis significantly increased both the amplitude (Fig. 1J, 1K) and frequency (Fig. 1J, 1L) of mEPSCs, suggesting upregulation of postsynaptic excitatory receptor expression as well as enhanced presynaptic excitatory input. Unexpectedly, we also observed increases in both mIPSC amplitude and frequency following CYP treatment (Fig. 1M, 1O–P), though statistical significance was detected only in the cumulative probability distributions (Fig. 1O left, 1P left), not in the averaged values (Fig. 1O right, 1P right). These findings imply that both presynaptic inhibitory input and postsynaptic receptor responsiveness may also be enhanced in the context of cystitis. Taken together, these results indicate that CYP-induced cystitis leads to robust, cell-type-specific changes in both intrinsic excitability and synaptic transmission within local CeA microcircuits, implicating these plastic changes as potential contributors to the neural processing of bladder pain, voiding dysfunction and associated affective symptoms in BPS.

Although slice physiology experiments reveal adaptive changes occurring in a subset of neurons within the CeA, they do not provide information about the specific molecular cell types in the CeA that are potentially involved in the pathophysiology of cystitis. To delineate the cellular populations within the CeA, first we performed single-nucleus multiome-sequencing (snMultiome-seq, combinatorial snRNA-seq and snATAC-seq) on dissected CeA tissue (Fig. 2A, Fig. S3A). This allowed us to identify the gene expression and chromatin accessibility underlying diversity of the CeA cell-types.

**Figure 2.**
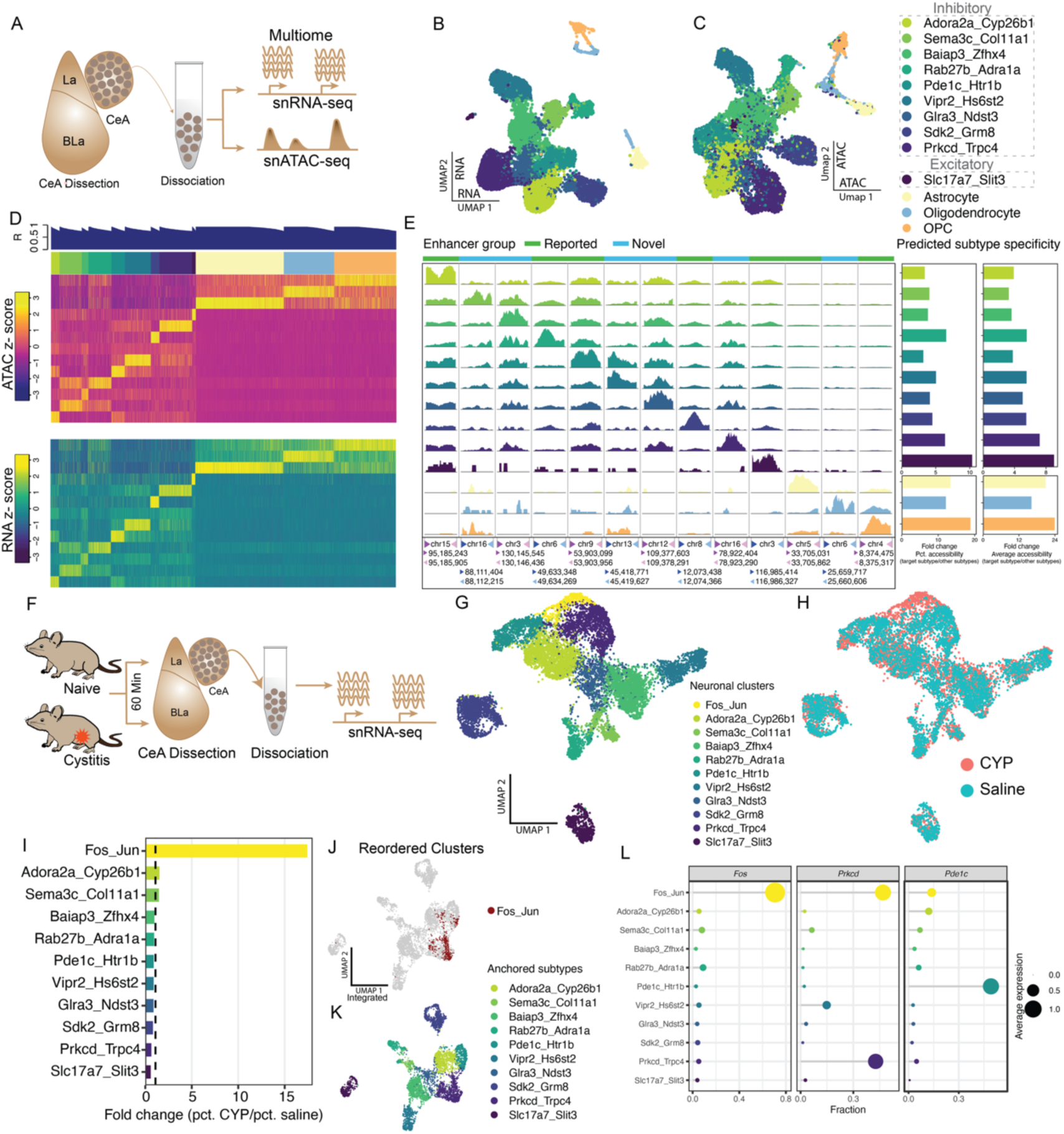
Single-nucleus multi-omic profiling of CeA reveals transcriptional cell states associated with CYP-induced cystitis. (A) Schematic of experimental workflow for CeA tissue microdissection to perform snRNA-seq and snATAC-seq. (B) UMAP visualization of snRNA-seq data shows transcriptional clustering of CeA cell types. (C) UMAP of snATAC-seq profiles reveals chromatin accessibility-based clustering of matched nuclei. Clusters were annotated with transcriptionally defined subtypes. (D) Heatmap show celltype-specific average expression of genes average accessibility of peaks in individual population. (E) Predicted enhancer regions grouped by previously reported and novel elements, with genomic coordinates and chromatin accessibility tracks across clusters. Bar plots (right) indicate the predicted subtype specificity of each enhancer. (F) Schematic for snRNAseq from naïve and cystitis mice. (G) UMAP of integrated saline and CYP snRNA-seq nuclei. CYP (pink) and saline (blue) cells are overlaid. (H) Cluster labels from integrated saline and CYP datasets, matching those from multiome data. (I) Bar graph showing fold change (CYP vs. saline) in cluster representation, identifying a marked expansion of the Fos_Jun–expressing cluster. (J) Re-embedded UMAP of neuronal nuclei, highlighting redistribution of Fos_Jun+ nuclei. (K) Neuronal cluster identity of Re-embedded UMAP. (L) Dot plots for Fos, Prkcd and Pde1c in neuronal subtypes.

We generated four snMultiome-seq libraries with no major batch variations and an average of 1,717.8 genes detected per nucleus for the snRNA-seq libraries and 6,900 high-quality transposase-sensitive fragments detected per nucleus for the snATAC-seq libraries (Fig. S3B,C). These transposase-sensitive fragments match expected nucleosomal size distribution (Figure S3D) and formed 116,506 peaks when aggregated across all snATAC-seq libraries. After quality control processes and removal of low-quality and doublet nuclei, we obtained 13,204 high-quality nuclei, including 12,158 neuronal nuclei and 1,046 non-neuronal nuclei. To identify the transcriptional cell types in the CeA, we first performed iterative clustering based on snRNA-seq data, which revealed 10 neuronal clusters (9 GABAergic types and 1 glutamatergic types) and 3 non-neuronal cell types (Fig. 2B), whose transcriptional profiles are in agreement with previous studies ^63–65^ and via Allen brain cell atlas (Figure S4).). Independent clustering of snATAC-seq data revealed that clusters identified based on chromatin accessibility closely mirrored transcriptionally defined cell types, with 91.4 ± 9.5% of nuclei in each epigenomic cluster are from the same transcriptional cell types, indicating a strong concordance between epigenomic and transcriptomic signatures within the CeA (Fig. 2C). To identify cell-type-specific genomic regulatory networks, we calculated the Pearson’s correlation between the accessibility of cell-type-specific peaks with the expression of cell-type-specific genes for any peak-gene pairs that are within a 5M bp window of the same chromosome. A peak-gene pair with a strong positive correlation (Pearson’s r > 0.75) is identified as a putative enhancer and its putative regulated gene. We identified 6,091 putative enhancer-gene pairs that are cell-type-specific for the CeA (Fig. 2D). Several lines of evidence support the validity and regulatory relevance of the putative enhancers identified in our study. First, 84.51% of the putative enhancer peaks are mapped to the distal non-coding region of the genome (Fig. S3E). These peaks exhibited a greater overlap with the DNase hypersensitive sites (96.5%) detected in the adult mouse brain compared to randomly selected genomic regions (41.1%) of similar sizes and GC contents. Second, these putative enhancers showed significant alignment with genomic regulatory elements previously annotated by the ENCODE consortium, particularly those identified in the mouse brain as opposed to other organs (Fig. S3F). To prioritize putative enhancers that may be used to precisely target individual cell types, we identified the top candidate in each cell type that exhibited the best cell type specificity by computational prediction (Fig. 2E), offering potential tools for targeting CeA subpopulations in future functional studies ^66–68^.

We next investigated whether cystitis selectively recruits specific CeA neuronal subtypes. We performed snRNA-seq of CeA from saline- and CYP-treated mice and were analyzed to assess changes in subtype engagement (Fig. 2F). We generated four snRNA-seq libraries with an average of 1,530.2 genes detected per nucleus (Fig. S5A). The snRNA-seq data exhibited minimal batch variations, with nuclei from different libraries broadly overlapping on the UMAP space (Fig. S5B). After quality control, a total of 16,096 high-quality nuclei were retained, including 11,212 neuronal nuclei and 4,884 non-neuronal nuclei. Iterative clustering revealed 11 neuronal clusters and five non-neuronal cell types (Fig. 2G, Fig. S5C,D). Among the CeA neuronal nuclei, there are 10 clusters that transcriptionally match the neuronal cell types prior defined in our snMultiome-seq data (Fig. 2B), with the nuclei composition of each cluster being relatively equal across saline- and CYP-treated mice (Fig. 2G-I). However, we found a new transcriptional cluster, Jun-Fos, where the majority of nuclei (95.1%) were derived from CYP-treated mice (Fig. 2G, H). This cluster is characterized by robust expression of more than 15 activity-dependent immediate early genes (IEG, Fig. S5G), such as *Fos* and *Jun* (Fig. 2H, I), suggesting this cluster represents a transcriptional state of CeA neurons activated by CYP. To identify the original transcriptional identity of nuclei in this cluster, we mapped these activated nuclei from CYP-treated mice to all neuronal nuclei from saline-treated mice using anchor analysis. This revealed that CYP-activated nuclei were derived selectively from two distinct neuronal subtypes: Prkcd-Trpc4 neurons defined by higher expression of *Prkcd* and *Trpc4*, and Pde1c-Htr1b neurons defined by higher expression of *Pde1c* and *Htr1b* compared to other neuronal subtypes (Fig. 2J-K), indicating subtype-specific activation in response to cystitis. Quantification of *Fos*, *Prkcd*, and *Pde1c* co-expression within confirmed the subtype-specific nature of this response, with each gene enriched in this cluster (Fig. 2L). Together, these findings identify Prkcd-Trpc4 and Pde1c-Htr1b CeA neurons being selectively recruited following CYP-induced bladder inflammation, suggesting a potential role for these subtypes in mediating cystitis-associated affective and sensory pain components characteristic of bladder pain syndrome (BPS).

Given the observed activation of molecularly heterogeneous CeA neurons in response to cystitis, we sought to selectively access and characterize the functional role of these stimulus-responsive ensembles in visceral pain sensitization and associated affective behaviors in cystitis. To this end, we employed the “Targeted Recombination in Active Populations 2.0” (TRAP2) mouse line ^69^, which expresses tamoxifen-inducible CreERT2 under the control of the *Fos* promoter. In this system, administration of 4-hydroxytamoxifen (4-OHT) enables temporally restricted genetic tagging of neurons that are transcriptionally active during a defined window of stimulus exposure. By aligning 4-OHT delivery with peak neuronal activation during acute cystitis induced by CYP, we were able to label CeA neurons that are selectively engaged during visceral pain induced by cystitis-referred to as FosTRAP neurons. To validate the specificity and efficacy of our activity-dependent neuronal tagging approach, we crossed FosTRAP mice with a Cre-dependent tdTomato flox-stop reporter line ^70^. Mice received either cyclophosphamide (CYP) or saline injections, concurrently with intraperitoneal administration of 4-OHT, allowing for temporally restricted CreERT2-mediated recombination and permanent expression of tdTomato in Fos-expressing neurons during cystitis induction (Fig. 3A, B). In CYP-treated mice, we observed robust tdTomato expression throughout the central amygdala (CeA) (Fig. 3C-F, Fig. S6A), while saline-injected controls showed sparse labeling (Fig. 3E, Fig. S6B). This minimal labeling in controls likely reflects limited neuronal activation in response to procedural stimuli, such as the injection itself, and may capture a small subset of pain-or stress-sensitive neurons.

**Figure 3.**
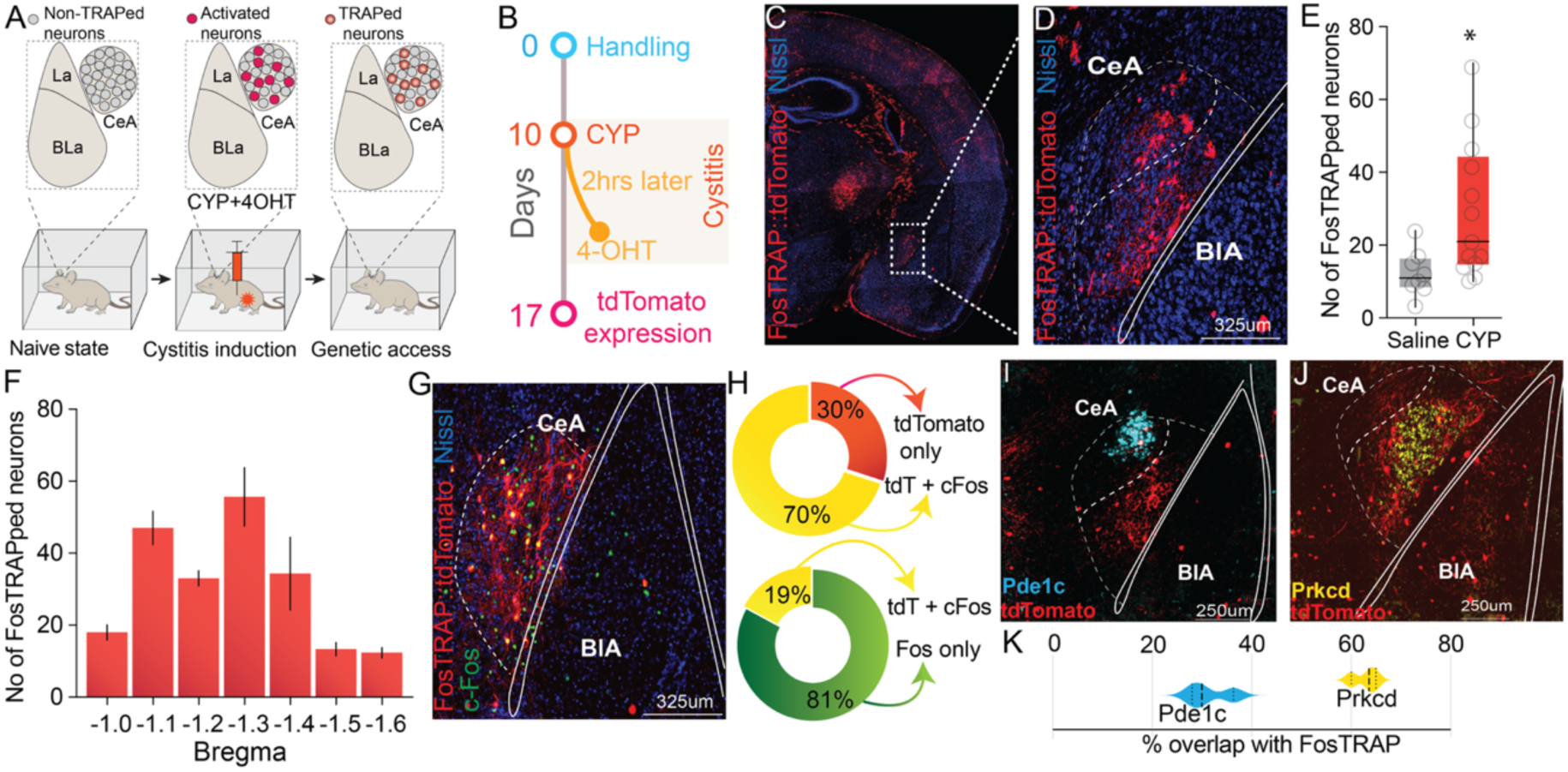
FosTRAPing of cystitis neuronal ensembles recruited in the CeA. **(A)** Schematic of FosTRAP strategy to selectively label cystitis-activated neurons in the CeA. (**B**) Timeline illustrating experimental TRAP strategy. **(C, D)** Representative immunofluorescence images showing cystitis activated FosTRAP+ neurons in CeA. Scale bars: 325μm. Dashed lines demarcate anatomical boundaries. **(E)** Quantification of FosTRAP+ neurons comparing saline vs CYP treatment groups (n=8-13 per group). *t test,* t=2.446, df=19, p = 0.024. **(F)** Analysis of FosTRAP+ neuron numbers across different bregma levels, showing peak targeting at −1.2 to −1.3 mm bregma. (**G**) Colocalization of CYP-activated cFos with tdTomato^+ve^ CYP FosTRAPed CeA neurons. **(H)** Pie chart showing relative percentages of Fos^+ve^ neurons that are tdTomato^+ve^ and tdTomato^+ve^ neurons that are Fos^+ve^. **(I-J)** Colocalization analysis of FosTRAP+ neurons with Pde1c and Prkcd markers. Scale bars: 250μm. **(K)** Quantification of Pde1c and Prkcd marker overlap with cystitis FosTRAP CeA neurons.

To assess the specificity and stability of FosTRAPed neurons in encoding cystitis-related activity (hereafter referred to as “cystitis-TRAPed neurons”), we re-exposed mice to a second CYP injection one week after the initial TRAPing event and performed immunohistochemistry for c-Fos protein 90 minutes later (Fig. 3G). A substantial proportion (70%) of tdTomato-positive CeA neurons co-expressed c-Fos in response to the second stimulus (Fig. 3H), demonstrating robust and reliable reactivation of the same neuronal ensemble upon re-challenge with CYP. These findings confirm that the FosTRAP strategy enables temporally precise and persistent genetic access to CeA neurons selectively activated during acute cystitis. This approach provides a powerful tool for dissecting the functional contributions of pain- and affect-related circuits in bladder pain syndrome (BPS).

As our single-cell studies revealed that acute cystitis induces robust upregulation of activity-dependent genes such as Fos specifically in in *Prkcd-Trpc4* and *Pde1c-Htr1b* CeA neuronal populations, we next sought to validate these findings. To this end, we performed RNAscope in situ hybridization for *Pde1c* and *Prkcd* in cystitis-TRAPed brains to assess their molecular identity within the activated ensemble (Fig. 3I–J).

Consistent with our transcriptomic data, we observed substantial colocalization between tdTomato+ cystitis-TRAPed neurons and these subtype markers. Quantitative analysis revealed that approximately 35% of tdTomato cells co-label *Pde1c+* and 65% of tdTomato cells co-label *Prkcd+* (Fig. 3K), indicating that these cell types are preferentially engaged during cystitis and reliably captured by the TRAP paradigm. These results validate the specificity and utility of the FosTRAP approach for accessing functionally relevant, stimulus-activated CeA ensembles in vivo. Importantly, they also confirm that the activated neuronal population is molecularly heterogeneous, composed primarily of transcriptionally defined GABAergic subtypes, *Pde1c+* and *Prkcd+* that were selectively activated in our single-nucleus transcriptomic analysis. This convergence between molecular and functional data underscores the critical role of these CeA subtypes in encoding the acute affective and sensory consequences of bladder inflammation.

Next, we sought to determine whether the cystitis-TRAPed CeA ensembles recruited during cystitis represent a modality-specific population, or whether these neurons are broadly responsive to other salient aversive sensory stimuli, such as itch or inflammatory somatic pain—two sensory modalities also processed within the CeA ^42,71–78^. To address this, one week after CYP-induced TRAPing, we administered either complete Freund’s adjuvant (CFA; a inflammatory agent, injected into the hind paw to cause somatic pain) or chloroquine (a pruritogen, injected into the nape of the neck) in independent cohorts of FosTRAPed mice. Ninety minutes following stimulation, brains were collected and immunostained for c-Fos to visualize neuronal activation in response to these distinct modalities. As a control, we also quantified Fos expression in home-cage conditions without any additional stimulation. Notably, exposure to CFA-induced paw inflammation (somatic pain) yielded a substantially higher degree of overlap (∼40%) with the cystitis-TRAPed population, indicating small portion of same CeA neurons recruited by visceral and somatic inflammatory pain stimuli (Fig. S6C, D). In contrast, fos-expressing neurons by chloroquine-induced itch and home-cage conditions resulted in ∼30% overlap with the cystitis-TRAPed CeA ensemble, suggesting that this level of colocalization may reflect baseline FosTRAP tagging in the CeA (Fig. SE-H). Together, these data suggest that the CeA ensemble activated by cystitis exhibits functional selectivity for visceral nociceptive processing, with a partial convergence with somatic pain circuits than with pruriceptive pathways.

To determine whether cystitis-TRAPed CeA neurons are functionally involved in modulating visceral pain and affective behaviors, we employed an optogenetic approach to manipulate the activity of cystitis-TRAPed neurons in a temporally precise and cell-type-specific manner. FosTRAP mice were injected with a Cre-dependent viral vector expressing the excitatory opsin Channelrhodopsin-2 (ChR2; AAV5-EF1a-DIO-ChR2-eYFP) or a control virus (AAV5-EF1a-DIO-eYFP) into the right CeA, and TRAPed during CYP-induced cystitis (Fig. 4A-C). Four weeks later, after the resolution of acute behavioral sensitization (Fig. S1B, C), we optically activated the TRAPed CeA neurons to assess whether their activation is sufficient to produce visceral pain, bladder voiding and affective-related phenotypes. Optical reactivation of cystitis-TRAPed CeA neurons significantly increased referred abdominal hypersensitivity in response to abdominal von Frey mechanical stimulation, indicating that these neurons are sufficient to evoke referred visceral hypersensitivity even after the original inflammatory insult has subsided (Fig. 4D). In contrast, no change in mechanical sensitivity was observed following optical stimulation in control mice expressing eYFP. Given the observed overlap between visceral and somatic pain ensembles in the CeA, we next evaluated whether reactivation of cystitis-TRAPed neurons altered sensitivity to somatic pain stimuli. Indeed, optical stimulation of ChR2-expressing CeA neurons significantly increased both mechanical and thermal sensitivity of the hind paw, consistent with a role for these neurons in modulating both visceral and somatic sensitivity (Fig. 4E, F). To evaluate whether activation of cystitis-TRAPed CeA neurons alters bladder voiding behaviors, we performed a voiding spot assay (VSA) using filter paper under UV detection (Fig. 4G). One month after CYP-induced cystitis and TRAPing, ChR2- and eYFP-expressing FosTRAP mice underwent optogenetic stimulation of CeA neurons during the voiding assay. Activation of ChR2-expressing TRAPed neurons resulted in a significant increase in the number of small-volume urine spots compared to eYFP controls (Fig. 4H, I). These changes indicate a shift toward increased urinary frequency and decreased voiding efficiency, characteristic of bladder hypersensitivity. These findings suggest that the reactivation of cystitis-associated CeA ensembles is sufficient to induce persistent alterations in bladder function, even in the absence of ongoing inflammation.

**Figure 4.**
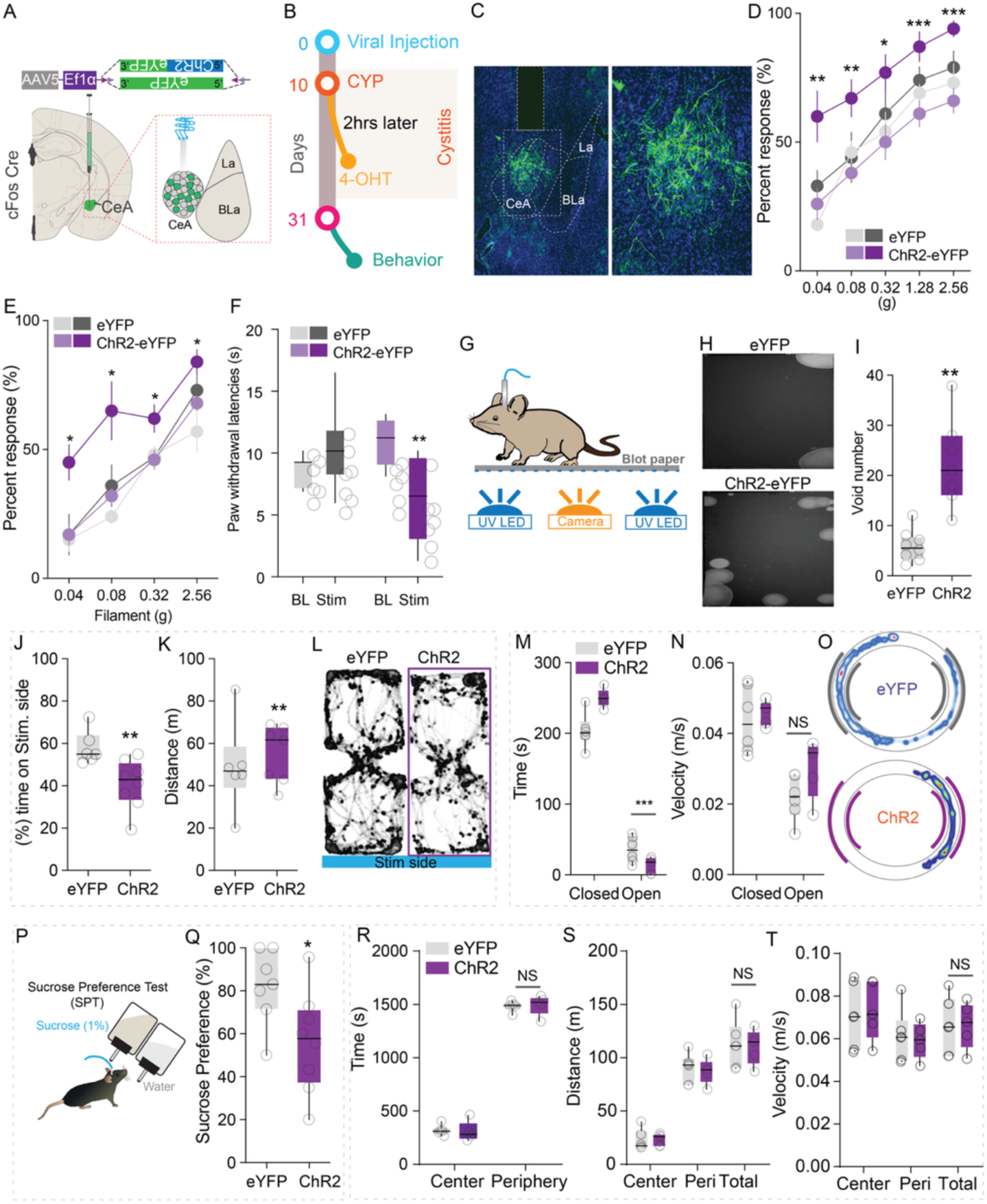
Optogenetic activation of FosTRAP-targeted neuronal ensembles causes referred pain, aversion, anxiety and anhedonia. **(A)** Experimental design schematic showing viral injection strategy with AAV-EF1α-DIO-ChR2 and eYFP constructs in CeA. **(B)** Timeline of CYP TRAP with viral injection and behavioral testing protocol. **(C)** Representative image of ChR2-eYFP expression in CeA. Scale bar: 325μm. **(D)** Abdominal, paw (**E**) von Frey responses and (**F**) thermal paw withdrawal latencies from baseline and following optogenetic stimulation in eYFP control vs ChR2-eYFP mice. n = 6–9 per group, * p < 0.05, **p < 0.01, ***P < 0.001, ANOVA and Bonferroni’s for post hoc tests. **(G-I)** Behavioral analysis setup for void spot assay. **(H)** Representative void spot blot papers from eYFP and ChR2 expressing mice following stimulation. (I) Void spot numbers from eYFP and ChR2 expressing mice following stimulation. n = 7–8 per group, *t test,* t t=4.942, df=13, p = 0.0003. (**J**) Total time spent and (**K**) distance traveled in the photostimulation-paired chamber for ChR2 and eYFP mice. n = 6-8 per group, t test, t = 3.062, df = 12, p=0.0099. (**L**) Representative heat maps of real-time place aversion assay with spatial location of ChR2 and eYFP mice during closed loop optical stimulation of FosTRAPed neurons. **(M)** Time spent in the open and closed arms, (N) Velocity following optogenetic activation of CeA TRAPed neurons in ChR2 and eYFP mice. Light stimulation was delivered entire time mice were on EZM. n = 10 per group. t test, t = 2.309, df = 18, p= 0.0330. (**O**) Representative heat maps of EZM assay with spatial location of ChR2 and eYFP mice. **(P)** Schematic of Sucrose preference testing. (**Q**) Percent sucrose preference of ChR2 and eYFP mice following optogenetic activation of FosTRAPed CeA neurons. Mann-Whitney test, n = 7-8 per group, p = 0.0389. (**R**) Time spent, (**S**) distance travelled, (**T**) velocity in ChR2 and eYFP mice following optical activation of FosTRAPed CeA neurons., n = 6–10 per group. Two-way ANOVA, p=0.9988 (time), p=0.2811(velocity), p=0.1810.

To investigate whether cystitis-TRAPed CeA neurons also encode affective dimensions of pain, we performed real-time place aversion (RTPA) testing. In this assay, animals were allowed to freely explore a two-chamber arena, with photostimulation of ChR2+ TRAPed neurons delivered selectively in one chamber. Mice with ChR2 expression in cystitis-TRAPed neurons exhibited robust avoidance of the stimulation-paired chamber, whereas control animals expressing eYFP showed no preference (Fig. 4J–L). This finding indicates that activation of these CeA neurons generates an internal state of negative valence, supporting their role in the affective component of visceral pain. Lastly, to assess whether cystitis-TRAPed neurons contribute to anxiety-like behaviors, we conducted elevated zero maze (EZM) testing. Reactivation of cystitis-TRAPed CeA neurons significantly reduced time spent in the open arms of the maze compared to controls (Fig. 4M–O), suggesting induction of an anxiogenic-like state. Importantly, reactivation did not elicit overt defensive behaviors such as freezing or escape in the open field test (Fig. 4R–T), indicating that these neurons selectively mediate anxiety-like responses without triggering fear-associated behaviors To assess whether cystitis-TRAPed CeA neurons also affect motivational states, we next performed a sucrose preference test (SPT), a classical assay for measuring anhedonia (Fig. 4P). Mice were given free access to water and 1% sucrose solution and received optical stimulation in ChR2- and eYFP-expressing FosTRAP mice. Optogenetic reactivation of ChR2-expressing cystitis-TRAPed neurons significantly reduced sucrose preference compared to eYFP controls, (Fig. 4Q), indicating a selective impairment in reward-related behavior. These results support a role for CeA cystitis-activated neurons in mediating negative affect and motivational deficits associated with chronic bladder pain.

To determine whether cystitis-TRAPed CeA neurons are necessary for the maintenance of bladder hypersensitivity and associated affective symptoms, we employed a loss-of-function strategy using the inhibitory opsin parapinopsin (PPO) ^79^. FosTRAP mice received AAV5-EF1a-DIO-PPO-Venus or a control AAV5-EF1a-DIO-eYFP virus targeted to the right CeA and were TRAPed during CYP-induced cystitis (Fig. 5A-C). Behavioral testing commenced six weeks post-TRAP, a time point when CYP-induced visceral hypersensitivity had resolved. During these assays, pulsed blue light was delivered to selectively silence the previously TRAPed CeA neurons. Initial optical inhibition of cystitis-TRAPed neurons in the absence of active inflammation did not affect abdominal mechanical sensitivity compared to pre-stimulation baselines or eYFP-expressing controls (Fig. 5D). We reasoned that this lack of effect may reflect the resolution of CYP-induced cystitis at this delayed time point (Supplementary Fig. S1A-C). To re-establish a state of bladder hypersensitivity, we administered a second CYP injection, which lead to spontaneous pain behaviors directed to the abdomen (licking, squashing), optical inhibition of the cystitis-TRAPed neurons significantly decreased the visceral pain behaviors in PPO expressing mice compared to the eYFP control mice (Fig. 5E, F). Next, to evaluate whether inhibition of cystitis-TRAPed CeA neurons is sufficient to ameliorate hyperactive bladder-like voiding behaviors observed due to cystitis reinstatement, we performed optical inhibition of the cystitis-TRAPed CeA neurons and performed the voiding spot assay. PPO-mediated inhibition of CeA TRAPed neurons resulted in a marked normalization of voiding patterns. Compared to baseline, optogenetic silencing of the cystitis-TRAPed neurons led to a significant reduction in the number of small, scattered urine spots, indicating improved bladder control (Fig. 5G, H). These parameters were not significantly altered in eYFP-expressing controls. These findings demonstrate that cystitis-TRAPed CeA ensembles contribute to bladder hypersensitivity and their acute silencing is sufficient to restore normal voiding behavior. Cystitis reinstatement reliably induced robust visceral mechanical hyperalgesia (Fig. 5I). Under this re-sensitized condition, optogenetic silencing of cystitis-TRAPed CeA neurons significantly attenuated referred hyperalgesia to von Frey stimulation of the abdomen relative to both baseline and eYFP controls (Fig. 5I). Notably, this suppression of pain sensitivity persisted for up to two hours following cessation of optical inhibition (Fig 5J), suggesting a sustained impact on downstream pain processing circuits (Fig. S7).

**Figure 5.**
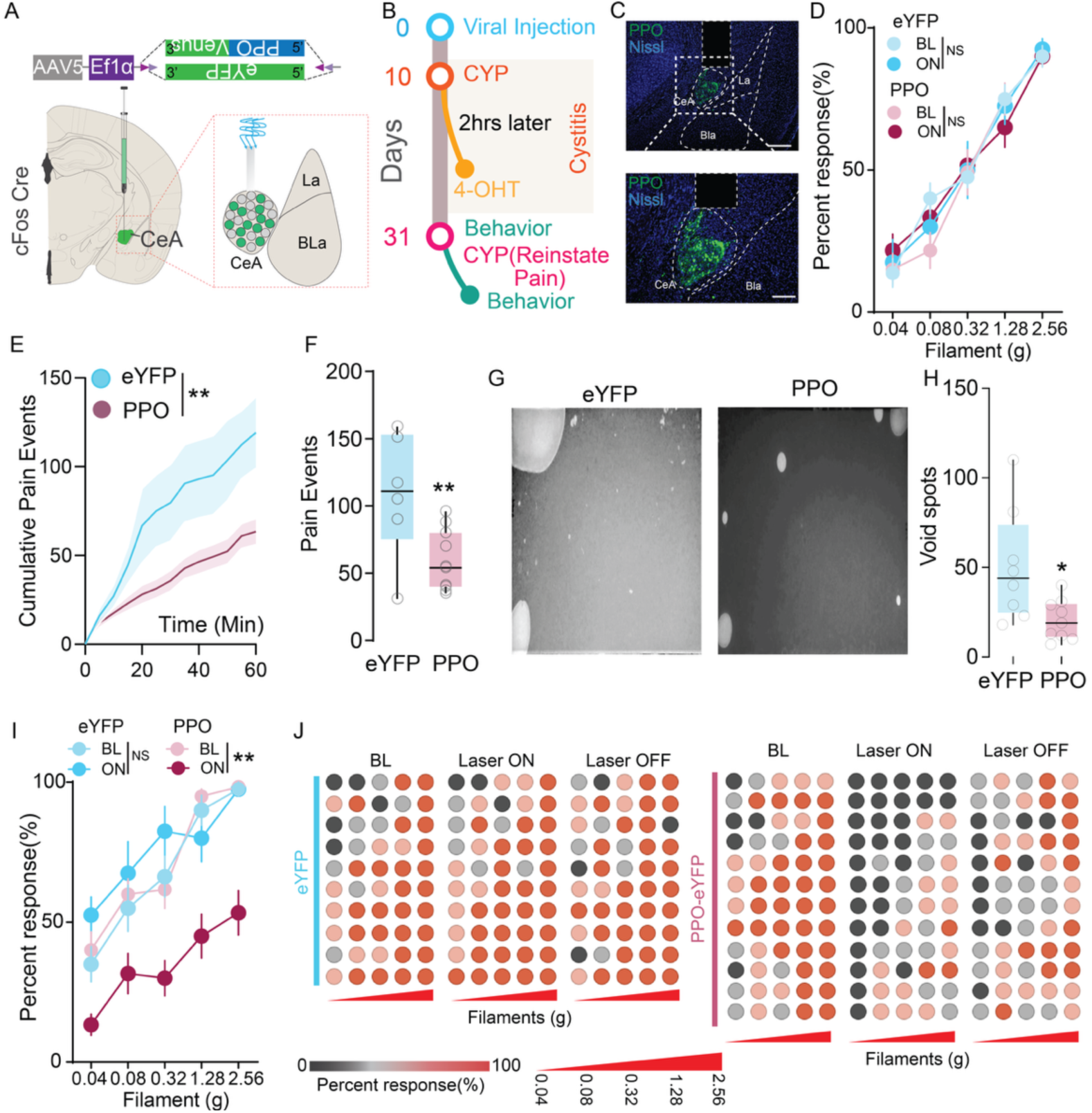
Inhibition of FosTRAPed CeA neurons attenuates spontaneous pain, referred pain and reveres bladder function. **(A)** Viral construct schematic for Cre-dependent PPO and eYFP expression in the CeA. **(B)** Experimental timeline for cystitis FosTRAP protocol. **(C)** Representative fluorescence images showing PPO expression in the CeA. **(D)** Abdominal von Frey responses from baseline and following optogenetic inhibition in eYFP control vs PPO mice. n = 8–10 per group, Two-way ANOVA, p>0.05. (**E**) Cumulative abdominal licks elicited following CYP injection comparing eYFP controls vs PPO-expressing mice. **(F)** Total abdominal licks in eYFP and PPO mice following optical inhibition. n= 12/group, t-test, t=2.992, df=24, p = 0.0063. **(G)** Representative blot papers from eYFP and PPO mice. **(H)** Quantification of void spots following optogenetic silencing in eYFP and PPO mice. n= 8-9/group, t-test, t=2.668, df=15, p = 0.0176. **(I)** Abdominal von Frey responses from baseline and following optogenetic inhibition in eYFP control vs PPO mice that are reinstated with cystitis. N=8-12/group, ***p < 0.001 **(J)** Heat map representation of individual animal responses to abdominal von Frey hair application from BL, laser ON and OFF conditions in eYFP and PPO-expressing animals.

To evaluate whether cystitis-TRAPed CeA neurons contribute to the affective dimension of bladder pain, we conducted a conditioned place preference (CPP) assay. Mice were allowed to freely explore a three-chamber apparatus, with mice were conditioned to optogenetic silencing of cystitis-TRAPed CeA neurons via parapinopsin restricted to one chamber (Fig 6A). In contrast to prior ChR2-mediated activation experiments, which induced robust avoidance of the stimulation-paired chamber, silencing of cystitis-TRAPed neurons elicited a significant preference for the light-paired side (Fig. 6B, C). This conditioned place preference indicates that inhibition of these neurons is positively reinforcing, likely due to attenuation of ongoing bladder pain and/or suppression of the negative affective state associated with cystitis. To further investigate the role of these neurons in anxiety-related behavior, we performed the elevated zero maze (EZM) assay (Fig 6D). Inhibition of cystitis-TRAPed CeA neurons resulted in a significant increase in both the time spent in the open arms and the number of open-arm entries compared to eYFP-expressing controls (Fig. 6E-G), consistent with an anxiolytic effect. These results suggest that cystitis-activated CeA ensembles are not only essential mediators of visceral pain but also contribute to anxiety-like behaviors, implicating a shared CeA circuit in the modulation of both nociceptive and affective components of cystitis. Finally, we confirmed that silencing cystitis-TRAPed CeA neurons did not produce generalized locomotor deficits or sedation. In the open field test, there were no significant changes in total distance traveled or velocity, and no freezing behavior (Fig. 6H, J), indicating that behavioral effects were not confounded by motor suppression.

**Figure 6.**
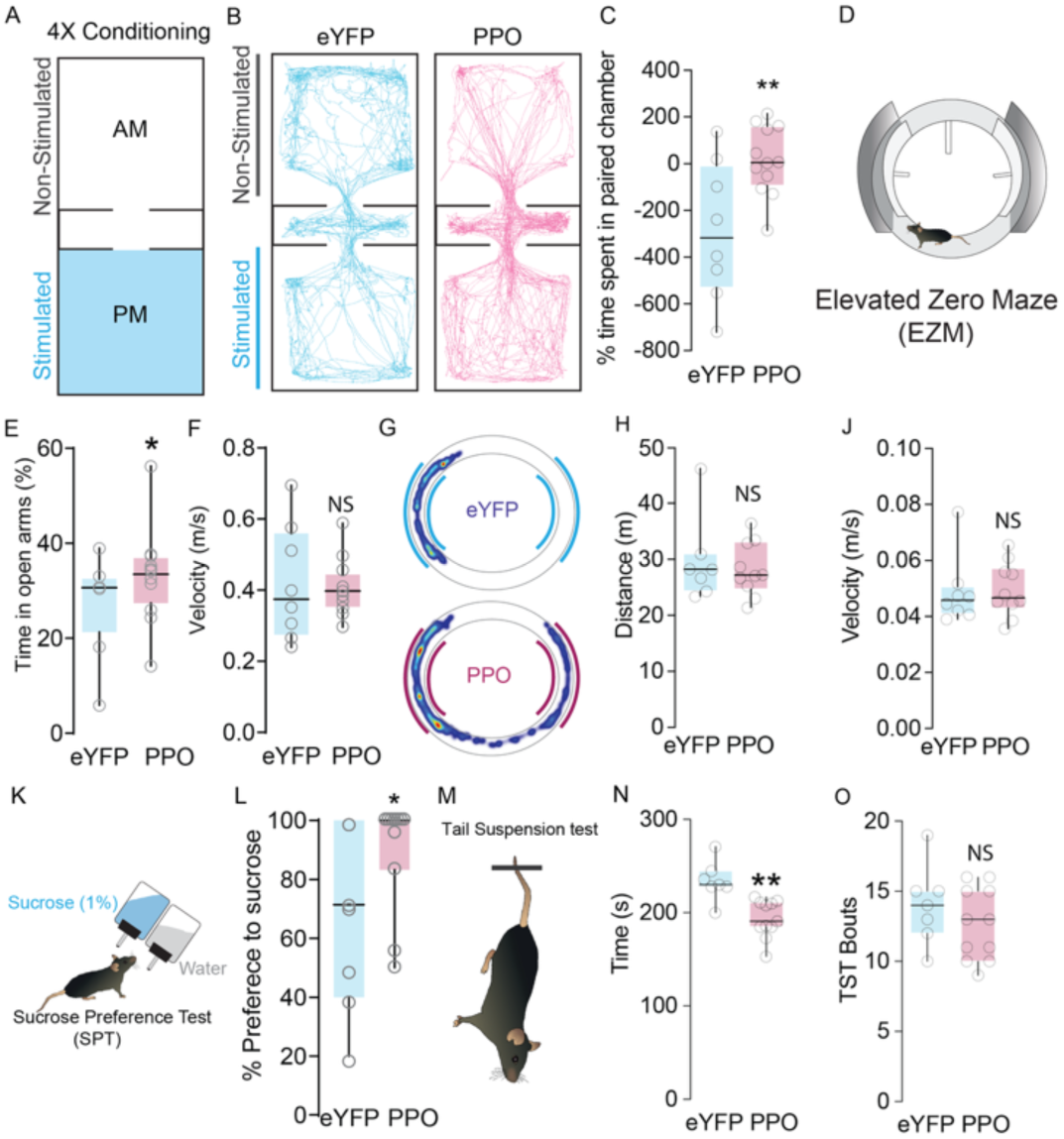
Inhibition of FosTRAPed CeA neurons attenuates aversive state, anxiety and anhedonia. (A) Conditioning paradigm schematic showing morning (AM) vs evening (PM) training sessions with eYFP and PPO group comparisons. (B) Representative movement tracks on post condition free run day, showing spatial location of eYFP and PPO injected mice after conditioning. (C) Change in chamber occupancy time in the stimulation of PPO-paired chamber compared to the eYFP-paired chamber. n= 9-11/group, t-test, t=2.184, df=18, p = 0.042. (D) Elevated zero maze schematic showing experimental set up. (E) Optical inhibition of PPO expressing CeA neurons increased time in the open arms of EZM compared to eYFP, control mice. n= 8-11/group, t-test, t=1.804, df=17, p = 0.038 (F) Optical inhibition of PPO FosTRAPed CeA neurons did not have any significant effect on motor behavior compared to the eYFP, control mice. n= 8-11/group, p>0,05. (G) Representative EZM movement tracks of eYFP and PPO during the optical inhibition. Distance traveled, (p=0.6224). (H), velocity, (p=0.8876), n = 6–10 per group. t test, (J). (K) Schematic of Sucrose preference testing. (L) Optogenetic inhibition of FosTRAPed CeA neurons leads to a significant increase in sucrose preference compared to the eYFP mice. n= 8-11/group. Mann-Whitney test, p = 0.068. (M) Representative schematic of tail suspension test. (N) Optogenetic inhibition of FosTRAPed CeA neurons significantly decreased immobility bouts compared to the eYFP controls. TST bouts (O) were not significantly different between eYFP and PPO-expressing mice (NS = not significant), suggesting specific modulation of pain-like behaviors without affecting basal activity or depressive-like behavior. n= 7-11/group, t-test, t=1.05, df=16, p = 3.075n= 7-11/group, t-test, t=4.054., df=16, p = 0.0009.

To further interrogate the affective contributions of cystitis-activated CeA neurons, we assessed anhedonia and behavioral despair—two core features of negative affective states—using the sucrose preference test (SPT) and the tail suspension test (TST), respectively (Fig. 6K, M). In the SPT, PPO-mediated optical inhibition of cystitis-TRAPed CeA neurons led to a significant increase in sucrose preference compared to eYFP-expressing controls (Fig. 6L), suggesting a reversal of cystitis-associated anhedonia. This enhancement in hedonic behavior indicates that CeA ensembles recruited by cystitis contribute to the suppression of reward sensitivity, a hallmark of maladaptive pain and affective comorbidity. Similarly, in the TST, inhibition of cystitis-TRAPed CeA neurons produced a marked reduction in immobility time relative to eYFP controls (Fig. 6M, N, O), reflecting decreased behavioral despair. These results, together with findings from the elevated zero maze and conditioned place preference assays, demonstrate that cystitis-activated CeA ensembles are not only involved in mediating visceral pain and bladder function but also play a critical role in the affective-motivational dimensions of cystitis.

## Discussion

Our electrophysiological analysis revealed that CYP-induced cystitis selectively enhances the intrinsic excitability of a subpopulation of CeA neurons (the adapting neurons). Notably we also observed that adapting neurons exhibited potentiated excitatory synaptic drive. Thus, our recordings show significant increases in both amplitude and frequency of mEPSCs in CYP-treated mice, indicating stronger glutamatergic input. Such changes likely reflect presynaptic and postsynaptic enhancements of excitatory transmission. This finding aligns with other studies of CeA synaptic plasticity in pain: for example, recent reports show that inflammatory pain markedly strengthened upstream synaptic inputs (PbN, BLa) via postsynaptic mechanisms ^42,48,73,76,80–83^. Our earlier work showed that pharmacological activation of CeA mGluR5 (a metabotropic glutamate receptor) drives bladder pain sensitization by increasing CeA output ^36^. In our case, the potentiation of excitatory currents in adapting neurons would be expected to amplify CeA output. In the context of the CeA microcircuit, increased excitation of adapting neurons could tip the balance of the dense local inhibitory network, effectively disinhibiting CeA output. In contrast, mIPSCs were only modestly altered by cystitis, changes in both the amplitude and frequency were small. This suggests that the net effect of cystitis might be a shift toward excitation. Indeed, some prior studies in neuropathic pain have emphasized loss of GABAergic inhibition in the amygdala as a contributor to hyperexcitability ^76,81,82,84^. These findings highlight that only subpopulations of the CeA undergo adaptive changes following cystitis.

The identification of distinct neuronal subtypes within the CeA—including nine GABAergic and one glutamatergic population—is consistent with recent single-cell profiling efforts ^64,65,85–88^., but extends this work by mapping cell-type-specific genomic regulatory elements using integrated chromatin accessibility data. We identified over 18,000 putative enhancer–gene pairs, a majority of which showed strong correspondence with ENCODE brain enhancers, further supporting the biological and functional relevance of our GRE dataset. This scATAC dataset from the CeA will hopefully amplify further development of new viral tools that can be tailored to target the CeA neuronal circuits to deliver the different gene cargos that can drive expression of cellular reporters, actuators and silencers.

In the context of cystitis, our single cell experiments revealed that although the global composition of CeA cell types remained unchanged in the CeA of saline mice, acute bladder inflammation induced selective activity dependent gene upregulation—most notably Fos and Jun in cell type specific manner. These results provide a molecular taxonomy of CeA neurons and pinpoint precise neuronal subpopulations selectively recruited by acute cystitis. Our results show that cystitis activates discrete GABAergic subpopulations within the CeA, advancing a cellular-resolution framework for understanding how CeA limbic circuits are recruited in the cystitis. Classical immunohistochemical and bulk sequencing approaches have reported broad classes of CeA neurons—including *Prkcd*+, *Som+, Penk+* and *Crf*+ subtypes as functionally important in pain and affect ^40,71,87–99^. Our single-cell data from control and cystitis mice reveal that a subset of transcriptionally defined cell types are selectively activated in response to acute cystitis. Specifically, two GABAergic subtypes defined by expression of *Prkcd–Trpc4* and *Pde1c–Htr1b* were significantly enriched for IEG activation following CYP-induced cystitis. This selective IEG induction indicates that these subtypes are likely involved in encoding the initial cystitis induced alterations in the CeA. These findings suggest that acute cystitis does not reorganize CeA cell composition but instead induces rapid, cell-type-specific functional plasticity, reflected in activity-state markers rather than global transcriptional remodeling.

The recruitment of the *Prkcd–Trpc4* population is especially notable as prior work revealed that *Prkcd*+ neurons in the CeL are known to be critical cell-type in maladaptive pain and negative affective states ^40,71,87,89–96^. While the CeA literature recognizes the critical role for *Prkcd*+ CeA neurons in nociceptive processing (pain sensitization and analgesia), our data extend this functional attribution by demonstrating that within the broader *Prkcd*+ population, a transcriptionally discrete subgroup defined by Trpc4 also has *Cartpt* expression (Fig. S5G) specifically necessary for cystitis induced maladaptive pain, bladder function and affective comorbidities. Additionally, we have also identified a novel cell-type in the CeA defined by expression of *Pde1c* and *Htr1b;* this novel cluster has not yet been yet studied in the context of pain, analgesia or affect. These results show that the *Pde1c–Htr1b+* cell-type in the CeA is activated by cystitis, suggesting a potential role in gating mechanism within specific CeA neurons during visceral pain processing. The concurrent activation of *Prkcd–Trpc4* and *Pde1c–Htr1b* suggests a potential convergence of behavioral sensitization and affective dysfunction in this CeA ensemble. Notably, despite robust IEGs activation within CeA specific clusters, our analyses revealed no significant changes in gene expression or cell type proportions between CYP- and saline-treated animals. This observation suggests that cystitis does not reorganize CeA cell-type composition within the acute time frame examined. Rather, pain-induced acute plasticity appears to rely on rapid, dynamic activation of subpopulations (*Prkcd–Trpc4* and *Pde1c–Htr1b*). Future studies should investigate CeA transcriptional changes either using bulk or single cell RNA sequencing to identify CeA subpopulations that play a role in chronicity of cystitis and provide high-resolution evidence in highly cell-type selective.

Interestingly, we found using the TRAP2 system that we were able to capture cystitis-activated CeA ensembles that have a high degree of overlap with the *Prkcd* and *Pde1c* subtypes identified by RNA-seq. Functional dissection of cystitis-TRAPed CeA neurons shows they are both sufficient and necessary to evoke symptoms associated with cystitis and associated phenotypes, including referred hyperalgesia, urinary dysfunction, anxiety-like behavior, and anhedonia. Our studies in cystitis-TRAPed neurons links activity, molecular identity, and behavioral output to demonstrate the contribution of these ensembles in cystitis symptomology. While optical silencing of the cystitis-TRAPed ensemble robustly attenuated both bladder hypersensitivity and negative affective behaviors, the specific functional contributions of each transcriptionally defined subpopulation remain to be clarified. It is plausible that distinct CeA subtypes within the broader TRAPed ensemble mediate separable dimensions of pain processing—e.g., one subpopulation of neurons encoding affective dysphoria, and another subpopulation contributing to visceral sensory gain or gating. Additionally, molecular markers we have identified could potentially label the adapting neurons we have shown with heightened intrinsic excitability observed in the cystitis model. Future work should study selective plasticity in *Prkcd–Trpc4+* and *Pde1c–Htr1b+* populations in cystitis using Cre lines or integrating our GRE dataset with enhancer-based AAV vectors could enable selective targeting of these pain-recruited CeA subtypes in vivo, offering a new level of cell-type and activity-state specificity in dissecting limbic pain circuits.

The modality-selective reactivation of cystitis-TRAPed neurons by somatic inflammatory stimuli but not by itch suggests shared convergence of somatic and visceral pain circuits within the CeA. In contrast, limited reactivation by itch stimuli indicates pruriceptive inputs engage distinct CeA populations, as suggested by prior findings from our group and others ^71,100–103^. This partial shared convergence of somatic and visceral pain circuits possibly offers insights into the observed cross-sensitization between bladder pain and other pain syndromes (e.g., fibromyalgia), and could explain why patients with IC/BPS often report comorbid somatic pain ^104–106^. In summary, our study provides causal evidence that a discrete, molecularly defined cystitis-activated ensemble of CeA neurons serves as a functional substrate for the progression of transient bladder inflammatory signals into maladaptive emotional memory associated with persistent visceral pain, which is a hallmark of IC/BPS. These findings provide a mechanistic foundation for future circuit-level dissection of chronic visceral pain.

## Methods

### Animals

All experimental procedures were approved by the Institutional Animal Care and Use Committee (IACUC) of Washington University School of Medicine and conducted in accordance with the National Institutes of Health guidelines for the care and use of laboratory animals. Mice were housed under standard conditions with a 12-hour light/dark cycle (lights on at 6:00 a.m.) and provided ad libitum access to food and water. All animals were maintained on a C57BL/6J genetic background, with no more than five animals per cage. Female littermates between 8 and 12 weeks of age were used for all experiments. As cystitis predominantly effects female, therefore, to conserve resources and minimize animal use in this labor-intensive study, all subsequent experiments were performed exclusively in female mice. FosCreERT2 mice (B6.129(Cg)-Fostm1.1(cre/ERT2)Luo/J; stock #030323), Ai14-tdTomato reporter mice (B6.Cg-Gt(ROSA)26Sortm9(CAG-tdTomato)Hze/J; stock #007914), and C57BL/6J mice were obtained from The Jackson Laboratory and breeding colonies were maintained in-house. For behavioral studies, heterozygous female FosCreERT2 mice were used. For Fos immunostaining and single-nucleus RNA sequencing experiments, female C57BL/6J mice (Jackson Laboratory) were used. Littermates were randomly assigned to experimental conditions, and all behavioral and analytical assessments were performed by investigators blinded to genotype and treatment group.

### Viral constructs

Purified and concentrated adeno-associated viruses coding for Cre-dependent ChR2-eYFP (rAAV5-DIO-ChR2-eYFP; 4.8 x 10^12^ particles/ml, Lot number: AV4313Y and Lot date: 04/21/2017) and control eYFP (rAAV5-DIO-eYFP; 3.3 x 10^12^ particles/ml, Lot number: AV4310i and Lot date: 07/21/2016) was used to express in the FosCreERT2 mice. All vectors were packaged by the University of North Carolina Vector Core Facility. PPO-Venus (AAV5:Ef1α:DIO:PPO-Venus;1.3×10^13^ vg/mL) was produced in the WashU Hope Center Viral Vector Core. All vectors were aliquoted and stored in −80°C until use.

### Stereotaxic Surgeries

Mice were anesthetized using 1.5–2.0% isoflurane in an induction chamber with an isoflurane/air mixture. Upon achieving a surgical plane of anesthesia, animals were secured in a stereotaxic apparatus (David Kopf Instruments, Tujunga, CA), and anesthesia was maintained with 2.0% isoflurane. Body temperature was maintained throughout the procedure using a thermostatically controlled heating pad. Preoperative care included the application of sterile ophthalmic ointment to prevent corneal desiccation, subcutaneous administration of 1 mL sterile saline, and disinfection of the surgical site with iodine solution. A small midline dorsal incision was made to expose the skull. Viral injections targeting the central amygdala (CeA) were performed using the following stereotaxic coordinates: −1.24 mm anteroposterior (AP) from bregma, ±2.8 mm mediolateral (ML), and −4.5 mm dorsoventral (DV) from the skull surface. Injections (75–100 nL) were delivered via a 2.0 μL Hamilton syringe using a stereotaxically mounted syringe pump (Micro4, World Precision Instruments) at a rate of 1 μL/10 min. Following injection, the syringe was left in place for an additional 10 min to facilitate viral diffusion prior to slow retraction.

For fiber optic cannula implantation, custom implants were constructed from 100 μm core diameter optical fiber (0.22 NA, Thorlabs) housed in zirconia ferrules (Thorlabs). Cannulas (5 mm in length) were stereotaxically implanted into either the CeA or the ventrolateral periaqueductal gray (vlPAG) using the following coordinates: CeA, −1.24 mm AP, ±2.8 mm ML, −4.25 mm DV; vlPAG, −4.84 mm AP, ±0.5 mm ML, −2.7 mm DV. Cannulas were affixed to the skull with two stainless steel bone screws (CMA, #7431021) and secured with dental cement. Cranial incisions were closed with sutures and reinforced with veterinary tissue adhesive. A topical triple antibiotic ointment was applied to the incision site. Postoperative monitoring continued heating pad until animals fully recovered from anesthesia. Mice were given a minimum of 14 days for recovery before behavioral testing commenced. Animals with misplaced viral injections or cannula placements outside the target regions were excluded from analysis.

### Optogenetic Manipulations

For all behavioral experiments involving optogenetics, mice were acclimated to fiber optic tethering for five consecutive days prior to the onset of testing. Animals were habituated using lightweight patch cables (Doric Lenses) connected to a 475 nm laser (Shanghai Laser), enabling free movement during stimulation sessions. To minimize mechanical constraints, the patch cables were coupled to an optical commutator (Doric Lenses), allowing for unrestricted locomotion while maintaining consistent optical connectivity. In FosCreERT2 mice, photostimulation was delivered via a programmable Arduino microcontroller, which was interfaced with the laser system to administer light pulses at frequencies of 5, 10, 20, (pulse width: 5-10 ms; light intensity: ∼10 mW/mm² at the fiber tip).

### Activity-Dependent FosTRAP Labeling

A 20 mg/mL stock solution of 4-hydroxytamoxifen (4-OHT; Sigma, Cat# H6278-10MG) was first prepared by dissolving 10 mg of 4-OHT in 500 µL of 100% ethanol via vortexing followed by sonication. This solution was subsequently diluted 1:4 in pre-warmed (45°C) autoclaved corn oil to achieve a final concentration of 5 mg/mL. The mixture was sonicated until fully dissolved and centrifuged under vacuum for 10 minutes to remove residual ethanol. Female FosTRAP mice (FosCreER2+/−; Ai14+/− or viral-injected FosCreER2+/−) were singly housed and gently handled for 7–10 days prior to experimentation to minimize stress-induced c-fos activation. On the day of the experiment, mice received a single intraperitoneal injection of 4-OHT (20 mg/kg) in their home cage. Sixty minutes post-4-OHT administration, mice were injected CYP (100 mg/kg, i.p) to selectively TRAP neurons activated by acute cystitis. In FosCreER2+/−; Ai14+/− mice, robust tdTomato expression was observed one-week post-TRAP. In FosCreER2+/− mice receiving optogenetic viral vectors, TRAPed populations were robustly labeled four weeks post-TRAP.

### Spontaneous Behavior Following CYP-Induced Cystitis

Immediately following intraperitoneal (i.p.) injection of cyclophosphamide (CYP) or saline, mice were individually housed in transparent plastic cages without sawdust bedding to facilitate behavioral monitoring. Each animal’s spontaneous behavior was continuously video-recorded for 60 min, and activity was quantified offline from the recordings by a blinded observer. Spontaneous behavior was classified into the following categories: Immobility: Periods of complete inactivity or resting posture. pain-like behaviors: Includes stretching, abdominal contractions, licking or biting of the lower abdomen, and writhing. If more than one of these signs was observed during a single interval, the corresponding values were summed. A final cumulative behavior score was obtained by summing the scores across all time points.

### Abdominal Mechanical Sensitivity

Abdominal mechanical sensitivity was assessed by quantifying the number of withdrawal responses to calibrated von Frey filament applications (North Coast Medical, Inc., Gilroy, CA; 0.02, 0.08, 0.32, and 1.28 g) delivered to the lower abdomen similar to our earlier work^107^. Each filament was applied 10 times with at least 15 seconds between individual applications and a minimum of 5 minutes between different filament forces. Mice were habituated in individual compartments on a raised plastic mesh platform for at least one hour prior to testing. All experimenters were blinded to mouse genotype and treatment group.

### Mechanical and Thermal Paw Sensitivity Assessment

Mechanical Sensitivity: Mechanical sensitivity was assessed using calibrated von Frey filaments (North Coast Medical, Inc., Gilroy, CA; 0.02, 0.08, 0.32, and 1.28 g) applied to both hindpaws, following previously established protocols ^71,107,108^. Mice were acclimated for at least one hour in individual enclosures placed on an elevated plastic mesh screen prior to testing. Each filament was applied 10 times per paw, with a minimum interstimulus interval of 15 seconds and at least 5 minutes between each filament size. The number of paw withdrawals in response to each stimulus was recorded and used as an index of mechanical sensitivity. Thermal Sensitivity (Hargreaves Test): Thermal nociception was evaluated using the Hargreaves assay (IITC Life Science, Model 390) as previously described (Samineni et al., 2017a). Radiant heat was applied to the plantar surface of each hindpaw, and paw withdrawal latency was recorded. Three trials were conducted per hindpaw, and the mean of the six measurements (three per paw) was used for analysis. A minimum intertrial interval of 5 minutes was maintained to prevent sensitization.

### Open-Field Test (OFT)

To assess locomotor activity, mice were acclimated to the testing room in their home cages for 2 hours prior to the start of testing. Mice injected with AAV constructs expressing ChR2 in the CeA were individually placed into a square open-field arena (55 × 55 cm; Noldus) located in a sound-attenuated room. Mice were allowed to freely explore the arena for 30 minutes while their behavior was recorded. Total distance traveled and locomotor patterns were analyzed using Ethovision XT software (Noldus Information Technologies, Leesburg, VA), as previously described^71^.

### Elevated Zero Maze (EZM)

Anxiety-like behavior was evaluated under low-light conditions (∼20 lux) using an elevated zero maze (Stoelting Co., Wood Dale, IL) consisting of two closed (30 cm wall height) and two open (1.3 cm wall height) quadrants on a circular track (60 cm diameter), positioned 70 cm above the floor (Montana et al., 2011). Mice were habituated to the testing room for 1 hour prior to testing. For ChR2 or control FosTRAP (eYFP) mice, mice were connected to optical fibers and received continuous 20 Hz photostimulation (5 ms pulse width) throughout the 6-minute trial. Each mouse was placed at the boundary between an open and closed section at trial onset. Movement was video-recorded using an overhead camera (Floureon HD), and behavioral parameters—including time spent in open arms, number of entries, and distance traveled—were analyzed using Ethovision XT.

### Real-Time Place Aversion/preference (RTPA, RTPP)

Aversion to optogenetic stimulation was assessed in a custom two-chamber apparatus (52.5 × 25.5 × 25.5 cm) filled with a layer of corn cob bedding, as described previously ^71^. Mice were placed in the neutral center zone and allowed to freely explore for 20 minutes. Entry into the stimulation-paired chamber triggered continuous blue light photostimulation (473 nm; 5 ms pulse width; 5, 10, 20, or 30 Hz; ∼10 mW light power). Entry into the opposite chamber terminated stimulation. Chamber assignment was counterbalanced across animals. Behavioral data, including time spent in each compartment and movement heatmaps, were analyzed using Ethovision XT.

### Conditioned Place Preference

Conditioned place preference (CPP) was conducted using an unbiased, counterbalanced three-chamber apparatus as described previously^71^, with modifications for bladder pain and optogenetic inhibition. Each of the two conditioning chambers was distinguished by unique visual and tactile cues (e.g., vertical vs. horizontal black-and-white stripes, distinct flooring textures), while a neutral center chamber connected them. On the first day, mice were allowed free access to all three chambers for 20 minutes to assess initial chamber preference. Video tracking was performed using Ethovision XT (Noldus). Mice displaying strong unconditioned preference (>70% time in any chamber) were excluded from further analysis. The two conditioning chambers were randomly assigned for laser OFF and laser ON pairings in a counterbalanced manner. On Days 2 through 4, mice underwent two daily conditioning sessions (morning and afternoon, ≥4 hours apart), each lasting 30 minutes. Morning sessions are paired with laser OFF and afternoon sessions are paired with laser ON. Optogenetic silencing was delivered throughout the conditioning trial. Laser light (473nm 10 Hz, 10ms, 4-5mW) was delivered through a rotary joint via patch cables coupled to the implanted fibers. On Day 5, mice were again allowed free access to all three chambers for 30 minutes. The preference score was calculated by subtracting the time spent in the laser-paired (PPO inhibition) chamber during the pre-conditioning session from the time spent in the same chamber during the post-conditioning session. Preference Score = Time_post (Laser ON chamber) – Time_pre (Laser ON chamber).

### Void spot assay

Mice were habituated to the behavioral testing room 24hrs prior to testing. On experiment day, behavioral testing was conducted during the light phase under dim light (20 lux). Testing equipment was cleaned with 70% ethanol between trials. The behavioral arena consisted of a UV-opaque acrylic chambers with no bottom placed on top of 0.35 mm chromatography paper (Fisher Scientific 05-714-4), which rested on a clear glass platform. Behavior was recorded using two wide-angle cameras (Logitech HD Cameras 720P) positioned below the arena. The bottom camera captured from beneath the glass surface. Videos were streamed at 15 frames per second with a resolution of 640×360 pixels. Illumination was provided by two UV fluorescent tube lights (American DJ Black-24BLB) placed below the arena and enclosed by foil walls to ensure even lighting of the chromatography paper. Mice were acclimated for 30 min prior to start of the trial and were given 1mL saline during this waiting time. Experimental trial lasted 2 hrs. During the session, optogenetic activation and silencing was delivered throughout the experimental trial. Laser light (473nm 20 Hz for ChR2, 10 Hz for PPO, 10ms, 4-5mW) was delivered through a rotary joint via patch cables coupled to the implanted fibers. At the end of the experiment, we took an image of the filter paper and counted the urine spots.

### Sucrose Preference Test (SPT)

The SPT was conducted as previously described^109^. Mice were habituated to the behavioral testing room 48hrs prior to testing. One day prior to experiment we water deprived for 18hrs. Mice were singly housed, and experiment was conducted 2hrs. During habituation, mice were exposed to two bottles of drinking water for 48hrs, followed by two bottles containing 1% sucrose solution. On the experiment day, bottles were assigned randomly. Following habituation, mice were water-deprived for 18hrs. During the test phase, mice were provided access to one bottle of 1% sucrose and one bottle of water for 2 hrs. During the session, optogenetic activation and silencing was delivered throughout the experimental trial. Laser light (473nm 20 Hz for ChR2, 10 Hz for PPO, 10ms, 4-5mW) was delivered through a rotary joint via patch cables coupled to the implanted fibers. Sucrose preference was calculated as the ratio of sucrose consumption to total fluid consumption over the 2-hour test period. The sucrose preference was calculated as the average of consumption of sucrose/ total consumption of water and sucrose during the 2 hr.

### Tail Suspension Test (TST)

The tail suspension test was performed as previously described ^110^ with modifications to allow optogenetic silencing. Mice were habituated to the testing room for at least 1 hour prior to the experiment. Each mouse was suspended by the tail using adhesive tape affixed ∼1 cm from the tip and hung from a horizontal bar 50 cm above a padded surface. A custom-built suspension apparatus allowed for tethering of the optical patch cables during the assay without interfering with movement. Mice were subjected to a single 6-minute trial. During the session, optogenetic silencing was delivered throughout the experimental trial. Laser light (473nm 10 Hz, 10ms, 4-5mW) was delivered through a rotary joint via patch cables coupled to the implanted fibers.

### Immunohistochemistry

Adult mice were deeply anesthetized with a ketamine/xylazine cocktail and transcardially perfused with 20 mL of phosphate-buffered saline (PBS), followed by 20 mL of 4% paraformaldehyde (PFA; w/v) in PBS (4 °C). Brains were carefully extracted, post-fixed in 4% PFA overnight at 4 °C, and cryoprotected by immersion in 30% sucrose (w/v) in PBS for at least 48 hours. Tissue was embedded in OCT compound and frozen at –80 °C before sectioning. Coronal brain sections (30 μm thick) were cut using a cryostat and stored in PBS containing 0.4% sodium azide at 4 °C until staining. To assess cFos expression in the CeA following CYP, pruritic stimulation, mice received a subcutaneous injection of chloroquine (200 μg/50 μL) to the nape of the neck and complete Freund’s adjuvant (CFA)-induced inflammatory pain, we injected 20ul into the paw. Ninety minutes later, mice were perfused as described. To verify the specificity of TRAPed tdTomato+ neurons for cystitis stimuli and exclude non-specific labeling, a separate cohort of FosCreER::Ai14 mice received a second CYP injection one week after TRAPing, followed by perfusion 90 minutes later.

For immunostaining, free-floating sections were washed in PBS and then incubated in a blocking solution containing 5% normal goat serum (NGS) and 0.2% Triton X-100 in PBS for 1 hour at room temperature. Sections were then incubated overnight at 4 °C in blocking solution containing the following primary antibodies: anti-mCherry (rabbit polyclonal, Clontech, #632543; 1:1000), anti-GFP (chicken polyclonal, Aves, #A11122; 1:2000), and anti-phospho-cFos (rabbit monoclonal, Cell Signaling, Ser32 D82C12; 1:2000). The following day, sections were washed three times in PBS and incubated for 1 hour at room temperature with fluorophore-conjugated secondary antibodies (all at 1:500; Life Technologies): Alexa Fluor 488 donkey anti-rabbit IgG, Alexa Fluor 488 goat anti-rabbit, Alexa Fluor 555 goat anti-mouse, and Alexa Fluor 555 goat anti-rabbit. NeuroTrace 435/455 (ThermoFisher, 1:500) was included to label neuronal cell bodies. After three additional PBS washes, sections were mounted on glass slides with Vectashield HardSet Mounting Medium (Vector Laboratories, H-1400) and allowed to cure before imaging. Fluorescent images were acquired using a Leica epifluorescence microscope.

### Fluorescence in situ hybridization (FISH)

To collect CeA samples for FISH studies, animals were anesthetized with ketamine cocktail and perfused with DEPC-PBS followed by 4% DEPC-PFA. Brain tissues were dissected and incubated in 4% DEPC-PFA at 4 C for 6 - 8 hours. Tissues were then incubated in 30% sucrose in 1X DEPC-PBS at 4 C for 24 - 48 hours until they sank to the bottom of the tube. Tissues were then embedded in OCT and stored at −80 °C before slicing. Tissues were sliced coronally into sections with 30 µm thickness using a cryostat (Leica), and tissue slices were mounted on microscope slides. Slides were stored at - 20 °C for a least one hour before moving to −80 °C for long-term storage. FISH was performed using the RNAscope® 2.0 Fluorescent Multiple Kit v2 User Manual for Fresh Frozen Tissue (Advanced Cell Diagnostics, Inc.). CeA sections were fixed in 4% paraformaldehyde, dehydrated, and treated with protease IV solution for 30 minutes. Sections were then incubated with specific target probes for the following genes: cFos (mm-Fos, catalog #316921), Pde1c (mm-Pde1c, catalog #489011) Prkcd (mm-Prkcd, catalog #432001). After probe hybridization, sections underwent signal amplification (AMP1–4) followed by incubation with fluorescently labeled probes (Opal 470, Opal 570, Opal 670), which were specific for the channels associated with each probe. After hybridization, slides were counterstained with anti-mouse RFP (Cat # 200-301-379) followed by Alexa 550 secondary to label tdTomato and DAPI and mounted with Vectashield HardSet Mounting Medium (Vector Laboratories). Imaging was performed on a Leica TCS SPE confocal microscope, and image analysis was conducted using the Application Suite Advanced Fluorescence (LAS AF) software.

### Acute brain slice preparation

The condition of electrophysiological recording was referred to Feng lab at MIT^111^ with slight modification. Briefly, mice were deeply anesthetized by intraperitoneal injection of Ketamine cocktail (concentration: 42.8 mg/ml. 30-50 μl per mouse) and then transcardial perfusion was performed with 25-30 mL of carbogenated NMDG aCSF. After perfusion, the mice were decapitated and the brain was extracted from the skull within 1 minute and placed into the carbogenated NMDG aCSF solution. Cutting off the undesired brain region and saved the desired the brain region. Glue the desired brain tissue onto the mounting cylinder. Pouring a sufficient volume of molten 2% agarose directly over the tissue on the mounting cylinder until completely submerged. After the rapid condensation of agarose by clamping the tube with the cold chilling block for 5-10 seconds, the lipstick tube with embedded brain block is inserted into the receiver of the slicing platform of the Compresstome VF-200 (Precisionary Instruments). Quickly filled the slicing chamber with NMDG aCSF solution with continued carbogenation throughout the procedure. Coronal slices with 300 μm thickness were obtained for all electrophysiological recording. Transferred the slices into a pre-warmed carbogenated NMDG aCSF solution for 10 min at 32-34 ℃. After the initial recovery period, transfer the slices into a new holding chamber containing room-temperature HEPES holding aCSF under constant carbogenation. The slices were stored for 1 hour at room temperature before transferring into the recording chamber for the use.

### Whole-cell slice electrophysiology

Electrophysiological recordings from right central amygdala (CeA) neurons from either rostral or caudal CeA (from −0.71 to −1.79 mm relative to bregma) in acute brain slice were performed in whole-cell patch-clamp configuration. The criteria for identifying the CeA was based on the atlas of Paxinos and Franklin (Fourth edition). Briefly, slices with the CeA were transferred to the recording chamber (Scientific Instruments) and perfused continuously (2.0 mL/min) with ACSF (continuously aerated with 95% O_2_/5% CO_2_) driven by a Mini-Peristaltic Pump II (Harvard Apparatus). The recording temperature was controlled at 31-34 ℃ by a HCT-10 Temperature Controller (ALA Scientific Instruments Inc.) for all recordings in this study. The CeA neurons were visualized using differential interference contrast optics on an upright microscope (BX50WI, Olympus). The recordings were performed using a MultiClamp 700B amplifier (Axon Instruments, Molecular Devices). The thin-walled borosilicate glass with filament (O.D.: 1.5 mm, I.D.: 1.1 mm, 10 cm length, fire polished, Sutter Instrument) was used in this study for obtaining recording pipettes by a horizontal puller (P-97, Sutter Instrument).

For the action potential recordings, electrodes (2.8 to 5.0 MΩ) were filled with the internal solution containing the following (in mM): 120 K-gluconate, 5 NaCl, 2MgCl_2_, 0.1 CaCl_2_, 10 HEPES, 1.1 EGTA, 4 Na_2_ATP, 0.4 Na_2_GTP, 15 Na_2_Phosphocreatine and 0.05% biocytin (adjusted pH to ∼7.3 with HCl and osmolarity to ∼290 with sucrose). The extracellular recording solution ACSF composed of (in mM): 124 NaCl, 2.5 KCl, 1.25 NaH_2_PO_4_, 24 NaHCO_3_, 5 HEPES, 12.5 Glucose, 1 MgCl_2_, 2 CaCl_2_, with synaptic blockers: 10 μM NBQX (2,3-Dioxo-6-nitro-1,2,3,4-tetrahydrobenzo[*f*]quinoxaline-7-sulfonamide disodium salt, AMPA/kainite antagonist), 50 μM D-APV (D-(-)-2-Amino-5-phosphonopentanoic acid, NMDA glutamate site antagonist), 10 μM Bicuculline ([*S*-(*R*\*,*S*\*)]-5-(6,8-Dihydro-8-oxofuro[3,4-*e*]-1,3-benzodioxol-6-yl)-5,6,7,8-tetrahydro-6,6-dimethyl-1,3-dioxolo[4,5-*g*]isoquinolinium iodide, GABA_A_ antagonist), 100 μM Picrotoxin (1:1 mixture of picrotoxinin and picrotin, GABA_A_ antagonist). After establishing the whole-cell recording configuration, at least 3 mins to wait before recording to allow adequate equilibration between the internal solution and the cell interior. Pipette capacitance was compensated before recording. The series resistance (Rs) was monitored but not compensated. The Rs was less than or equal to 32.3 MΩ in the data set in this study. Current clamp bridge balance was adjusted prior to each AP family recording. Recordings were low-pass filtered at 3 kHz and sampled at 10 kHz. To assess spontaneous excitatory and inhibitory synaptic events, whole-cell voltage clamp was configurated. Electrodes (2.8 to 5.0 MΩ) were filled with pipette solution containing the following (in mM): 110 D-gluconic acid, 110 CsOH, 0.1 CaCl_2_, 4 MgATP, 0.4 Na_2_GTP, 10 Na_2_Phosphocreatine, 1.1 EGTA, 10 HEPES, 8 TEA^+^Cl^-^, 3 QX314^+^Br^-^. The pH and osmolarity of internal solution were adjusted to 7.3 and 290 mOsm/l, respectively. To record the miniature EPSCs (mEPSCs) or IPSCs (mIPSCs), 1 μM TTX was added to the extracellular recording solution to block the action potential conduction. For the mEPSCs, the extracellular recording solution ACSF composed of (in mM): 124 NaCl, 2.5 KCl, 1.25 NaH_2_PO_4_, 24 NaHCO_3_, 5 HEPES, 12.5 Glucose, 1 MgCl_2_, 2 CaCl_2_.

GABA receptor antagonists (10 μM Bicuculline + 100 μM Picrotoxin) were added to isolate EPSCs. For the mIPSCs, the extracellular recording solution ACSF composed of (in mM): 124 NaCl, 2.5 KCl, 1.25 NaH_2_PO_4_, 24 NaHCO_3_, 5 HEPES, 12.5 Glucose, 1 MgCl_2_, 2 CaCl_2_. Glutamate receptor antagonists (10 μM NBQX + 50 μM D-APV) were added to isolate IPSCs.

### Tissue Collection for snRNA-seq

To collect mouse CeA tissues for sequencing, animals were anesthetized with a ketamine/xylazine cocktail and transcardially perfused with ice-cold NMDG-based cutting solution (NMDG 93 mM, KCl 2.5 mM, NaH2PO4 1.25 mM, NaHCO3 30 mM, HEPES 20 mM, Glucose 25 mM, Ascorbic acid 5 mM, Thiourea 2 mM, Sodium Pyruvate 3 mM, MgSO4 10 mM, CaCl2 0.5 mM, N-acetylcysteine 12 mM; pH adjusted to 7.3 with 12N HCl, bubbled with 95% O2 and 5% CO2). The brain was rapidly removed and dissected while submerged in ice-cold NMDG-based cutting solution. Coronal brain slices (400 µm thick) were obtained using a Compresstome (Precisionary Instruments, VF-210-0Z). The CeA were then micro-dissected under a stereomicroscope (Leica S9i) in a sterile petri dish that had been treated with RNAse-X and kept on dry ice. A reusable 0.75 mm biopsy punch (WPI 504638) was used to isolate the CeA tissue. For each sequencing sample, bilateral CeA tissues from five mice were collected and pooled into a nuclease-free centrifuge tube, which was kept on dry ice throughout the procedure. Samples were immediately stored at −80°C until nuclei isolation for subsequent processing.

### Single-Nuclei Isolation

Nuclei extraction from mouse CeA tissues was performed using a density gradient centrifugation protocol, modified from a previously described method to accommodate smaller tissue volumes (85). Frozen tissue samples were transferred from −80 °C and briefly thawed on ice for 2 minutes. Tissues were then transferred into a Dounce homogenizer (Millipore Sigma, D8938-1SET) containing homogenization buffer (0.25 M sucrose, 25 mM KCl, 5 mM MgCl₂, 10 mM Tris-HCl, pH 8.0, 5 µg/mL actinomycin D, 1% BSA, 0.08 U/μL RNase inhibitor, and 0.01% NP-40) kept on ice. Homogenization was performed with 10 strokes of the loose pestle followed by 10 strokes with the tight pestle in a total volume of 1 mL. For snRNA-seq samples, homogenate was centrifuged at 500 × g for 10 minutes at 4°C, and nuclei pellet was resuspended in resuspension buffer (1× PBS with 1% BSA and 0.08 U/μL RNase inhibitor) supplemented with 5 ng/μL 7-AAD. A density gradient centrifugation was applied to snMultiome-seq samples to enrich for neuronal nuclei. Homogenate was mixed 1:1 with a working solution consisting of 50% iodixanol, 25 mM KCl, 5 mM MgCl₂, and 10 mM Tris-HCl (pH 8.0). The nuclei suspension was then layered onto an iodixanol gradient and subjected to ultracentrifugation. Following centrifugation, nuclei were collected from the interface between the 30% and 40% iodixanol layers and diluted in the resuspension buffer. Nuclei were pelleted at 500 × g for 10 minutes at 4°C, resuspended in resuspension buffer supplemented with 5 ng/μL 7-AAD. Nuclei were subjected to fluorescence-activated cell sorting (FACS) to eliminate cellular debris. 7-AAD-positive events were sorted using a 100 μm nozzle on a BD FACSAria III into sterile 1.5 mL microcentrifuge tubes.

### snRNA-seq and snMultiome-seq

Sorted nuclei were counted using a hemocytometer and diluted to the appropriate concentration to target the recovery of 10,000 nuclei per library. snRNA-seq libraries were prepared using the 10x Genomics Chromium Single Cell 3’ Gene Expression v3 platform according to the manufacturer’s instructions. For snMultiome-seq, nuclei were processed and sequenced according to the manufacturer’s manual of 10X Genomics Chromium Single Cell Multiome Assay. Libraries were sequenced on an Illumina NovaSeq 6000 system, with 150 cycles each for Read 1 and Read 2, targeting a depth of 50,000 paired-end reads per nucleus for snRNA-seq libraries and 25,000 pair-end reads per nucleus for sATAC-seq libraries. Raw sequencing data were processed using the Cell Ranger software suite (cellranger v6.1.2 or cellranger-arc v2.0.1, 10x Genomics). Reads were aligned to the mouse reference mm10.

### Quality Control and Clustering of snRNA-seq Data and snMultiome-seq

The processed data from cellranger outputs were analyzed using the R (v4.4.1) package Seurat (v4.3.0) and Signac (v1.13.0). Nuclei were filtered based on the following quality control criteria on snRNA-seq data: a minimum of 500 detected genes, fewer than 15,000 total unique molecular identifiers (UMIs), and no more than 5% mitochondrial transcript content, and snATAC-seq data: at least 1,000 fragments overlapping called peaks. For snRNA-seq data, gene expression counts were normalized using the NormalizeData() function, which scales each nucleus to 10,000 total transcripts to account for differences in sequencing depth. Data were then scaled and centered using the ScaleData() function. Highly variable genes were identified with FindVariableFeatures(), and dimensionality reduction was performed using principal component analysis (PCA) via RunPCA(), retaining the top 20 principal components (PCs). Uniform Manifold Approximation and Projection (UMAP) was employed for visualization using RunUMAP(), based on the top 20 PCs. Clustering was carried out using the FindClusters() function with a resolution parameter set to 0.6. Marker genes for each cluster were identified using FindAllMarkers() by comparing each cluster to all others, applying a false discovery rate (FDR) threshold of <0.05 and a log2 fold-change >1. Clusters were flagged as low-quality or potential doublets if they met either of the following criteria: (1) lack of significantly enriched marker genes (FDR < 0.05, log2FC > 1), or (2) presence of five or more mitochondrial genes among the top 20 markers (ranked by average log2 fold-change). Clusters were annotated as neuronal if they showed high expression of canonical neuronal markers (e.g., Rbfox3, Snap25, Syt1), or as non-neuronal if they expressed markers of glial or other cell types (e.g., Sparc, Mbp, Apoe). Neuronal and non-neuronal nuclei were further sub-clustered separately, and low-quality or doublet-enriched clusters were re-evaluated using the same criteria described above. For snATAC-seq data, normalization of accessibility profiles was performed using term frequency-inverse document frequency (TF-IDF) via the RunTFIDF() function. Variable features were selected using FindTopFeatures(), and dimensionality reduction was performed using singular value decomposition (SVD) via RunSVD(). Uniform Manifold Approximation and Projection (UMAP) embeddings were computed with RunUMAP().

### Differential analyses of gene expression and chromatin accessibility

To identify transcriptional cell-type-specific features, differential expression and accessibility analyses were conducted using the FindAllMarkers() function in Seurat/Signac, comparing nuclei from a given cell type or projection to all other nuclei. Genes or peaks with a false discovery rate (FDR) < 0.05 were considered significant. For pairwise comparisons between two specific subtypes or projection-defined populations, FindMarkers() was used, again with significance thresholds of FDR < 0.05. To identify the transcriptional response, including immediate early genes (IEGs), to CYP treatment in the Fos_Jun cluster, features with an average log₂ fold change (avg_log2FC) > 0.5 and FDR < 0.05 were reported.

### Identification of putative genomic regulatory elements

To identify cis-regulatory elements underlying cell type-gene regulation, we adapted a previously described pipeline ^112,113^. For each peak and gene, we calculated average chromatin accessibility (snATAC-seq) and expression levels (snRNA-seq) across either transcriptional subtypes. Peak-gene pairs were restricted to those located within 5 Mb of each other on the same chromosome (measured from peak center to transcription start site [TSS]). For each pair, Pearson correlation coefficients were computed to assess coordinated regulation. Putative cell type-specific GREs were identified by first detecting snATAC-seq peaks and snRNA-seq genes enriched in the same subtype (avg_log2FC > 0.5 and FDR < 0.05 versus all others). Pairs with strong positive correlations (Pearson’s r > 0.75) were designated putative enhancers (pu.Enhancers); those with strong negative correlations (r < −0.75) were labeled putative silencers (pu.Silencers).

### Statistical Analysis

All data analyses were conducted with experimenters blinded to the experimental conditions. Exclusion criteria included failure to localize expression in the experimental model or off-target virus/drug delivery. At least three replicates per condition were performed, and data were averaged for each behavioral assay. The number of animals used in each experiment is denoted by “N.” For statistical comparisons, normality was first assessed using the D’Agostino and Pearson omnibus normality test. Parametric tests, including paired t-tests and two-way ANOVA with Bonferroni post hoc corrections, were used when normality assumptions were met. If normality was not assumed, nonparametric tests (e.g., Wilcoxon matched-pairs test) were applied. Statistical significance was set at p < 0.05 for all tests.

## Data and code availability

Raw and processed data of snRNA-seq and snATAC-seq experiments included in this study are deposited to the NCBI Gene Expression (GEO) SRA with accession number GSE299703.

## Acknowledgements

This work was funded by NINDS R01NS106953 and NIDDK R01DK116178 to RWG, the Urology Care Foundation Research Scholars Program and Kailash Kedia Research Scholar Award and NIDDK Career Development award (K01 DK115634), NIDDK R01DK128475 to VKS. We thank the Flow Cytometry & Fluorescence Activated Cell Sorting Core in the Department of Pathology and Immunology at Washington University School of Medicine for help with FACS. We thank the Genome Technology Access Center at McDonnell Genome Institute at Washington University School of Medicine for help with snRNA-seq and snATAC-seq library construction and sequencing. The Center is partially supported by NCI Cancer Center Support Grant #P30 CA91842 to the Siteman Cancer Center from the National Center for Research Resources (NCRR), a component of the National Institutes of Health (NIH), and NIH Roadmap for Medical Research. We thank Michael Bruchas and Bryan Copits for sharing PPO virus. We would like to thank all the Samineni and Gereau lab members and WashU Pain Center colleagues for their help with manuscript preparation.

## Author contributions

V.K.S and R.W.G. designed the experiments; V.K.S., J.N.S, S.B.S. performed behavior, V.K.S., H.J.H, Y.Z. performed anatomical analyses; V.K.S., A.C, L.Y. performed library prep and FACS for RNA-seq; L.Y, B.Z, Performed data curation, analysis and methodology for RNA-seq; J.N.L. performed slice electrophysiology; V.K.S, J.N.S, L.Y, J.N.L, B.Z analyzed the data; V.K.S., and R.W.G. wrote the manuscript with comments from all the authors.

## Supplementary figures

**Figure S1. (Related to Figure 1).**
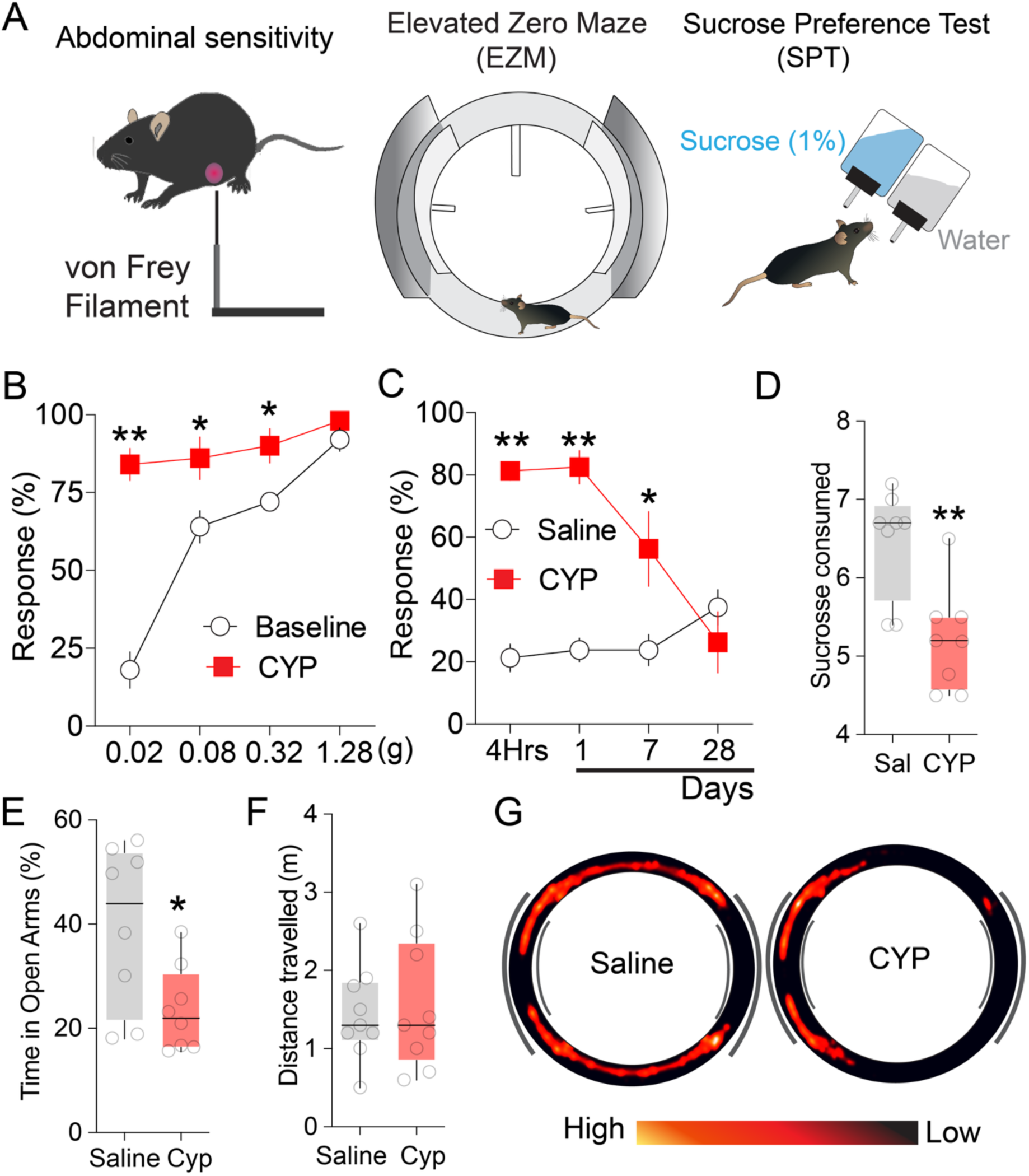
Cyclophosphamide induced mechanical sensitivity, anxiety and anhedonia symptoms. (A) Schematic illustrating behavioral models used in this grant proposal measuring abdominal sensitivity, Elevated Zero Maze, EZM, sucrose preference test, SPT. (B) Single dose of CYP (100 mg/kg) administration resulted in significant abdominal sensitivity to von Frey stimulation. (C) CYP induced abdominal sensitivity was observed starting 4hrs. post administration and lasted for one week. n=7 mice for each group, two-way repeated measures ANOVA. (D) One week post CYP, mice exhibited significantly decreased sucrose consumption compared to saline treated mice. n=8 mice for each group, two-tailed unpaired t test. (E) One week post CYP treatment, mice exhibited decreased time in the open arms of EZM compared to saline treated control mice, indicating development of anxious behavior. (F) CYP treatment did not have any significant effect on motor behavior compared to the saline treatment. (G) Representative heat maps show saline treated mice explore more in the EZM compared to the CYP treated mice. n=8 mice for each group. two-tailed unpaired t test, *p<0.05, **p<0.01.

**Figure S2. (Related to Figure 1).**
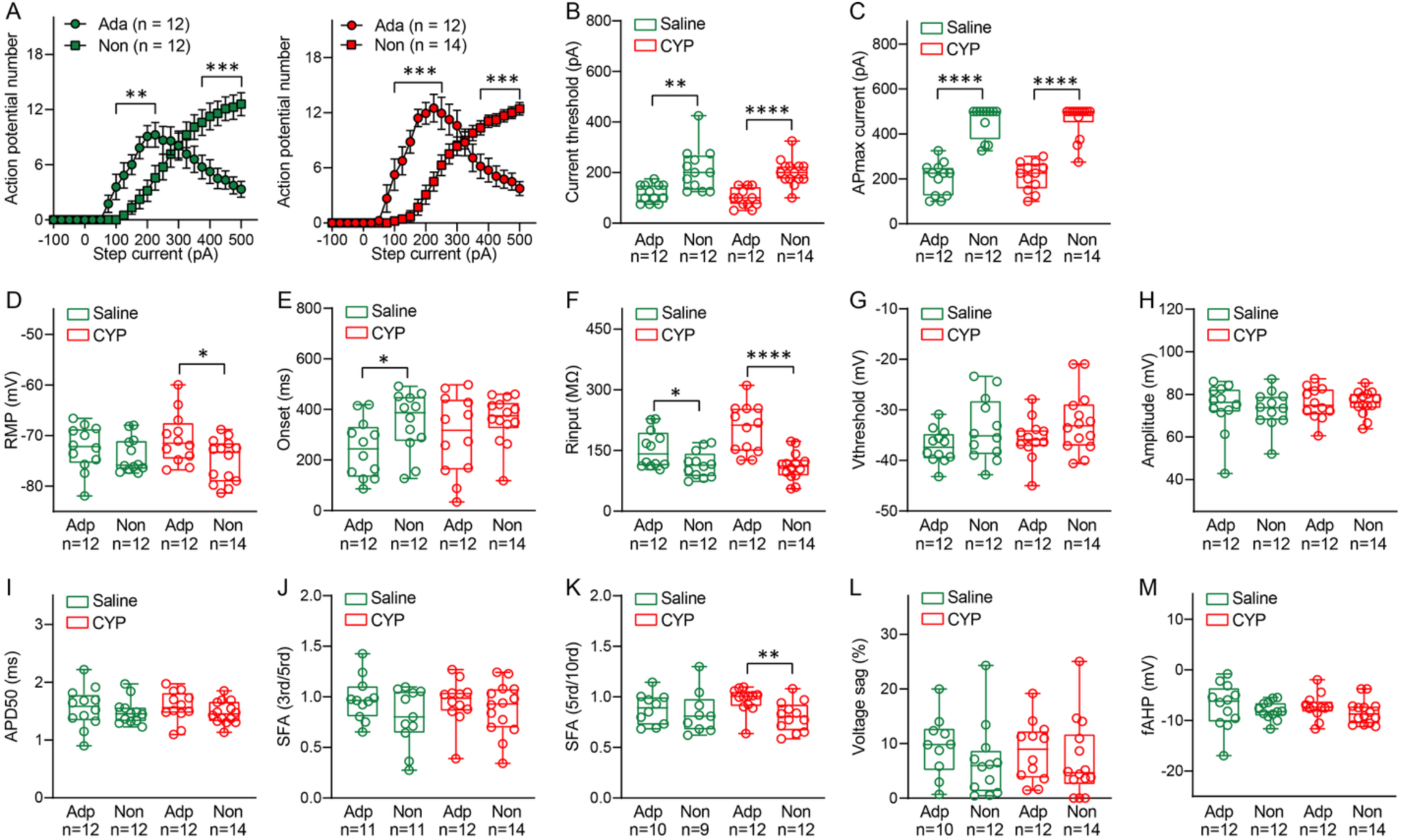
Two intrinsically distinct neural types in CeA showing strikingly different firing and subthreshold properties. (A) Plot showing comparison of AP number vs. current injection between adapting and non-adapting CeA neurons from saline-injected mice (left). Plot showing comparison of AP number vs. current injection between adapting and non-adapting CeA neurons from CYP-injected mice (right). (B) Boxplot displaying the comparison of current threshold between adapting and non-adapting CeA neurons from either saline-injected mice (Green) or CYP-injected mice (Red). (C) Boxplot displaying the comparison of current value evoking maximum AP number between adapting and non-adapting CeA neurons from either saline-injected mice (Green) or CYP-injected mice (Red). (D-M) Boxplots displaying the comparison of resting membrane potential (RMP), onset, input resistance (Rinput), voltage threshold for AP firing (Vthrehold), AP Amplitude, AP half-width (APD50), spike frequency adaptation (SFA(3rd/5rd)), slow spike frequency adaptation (SFA(5rd/10rd)), voltage sag and fast afterhypolarization potential (fAHP) between adapting and non-adapting CeA neurons from either saline-injected mice (Green) or CYP-injected mice (Red). Two-way ANOVA followed by Bonferroni multiple comparisons test in (A). Two-tailed unpaired t test for (B, red), (D), (E), (F, red), (G), (H), (1), (J), (K) and (M, red). Two-tailed Mann-Whitney test for (B, green), (C), (F, green), (-) and (M, green). * p < 0.05, **p < 0.01, ***P < 0.001, ****P < 0.0001.

**Figure S3. (Related to Figure 2).**
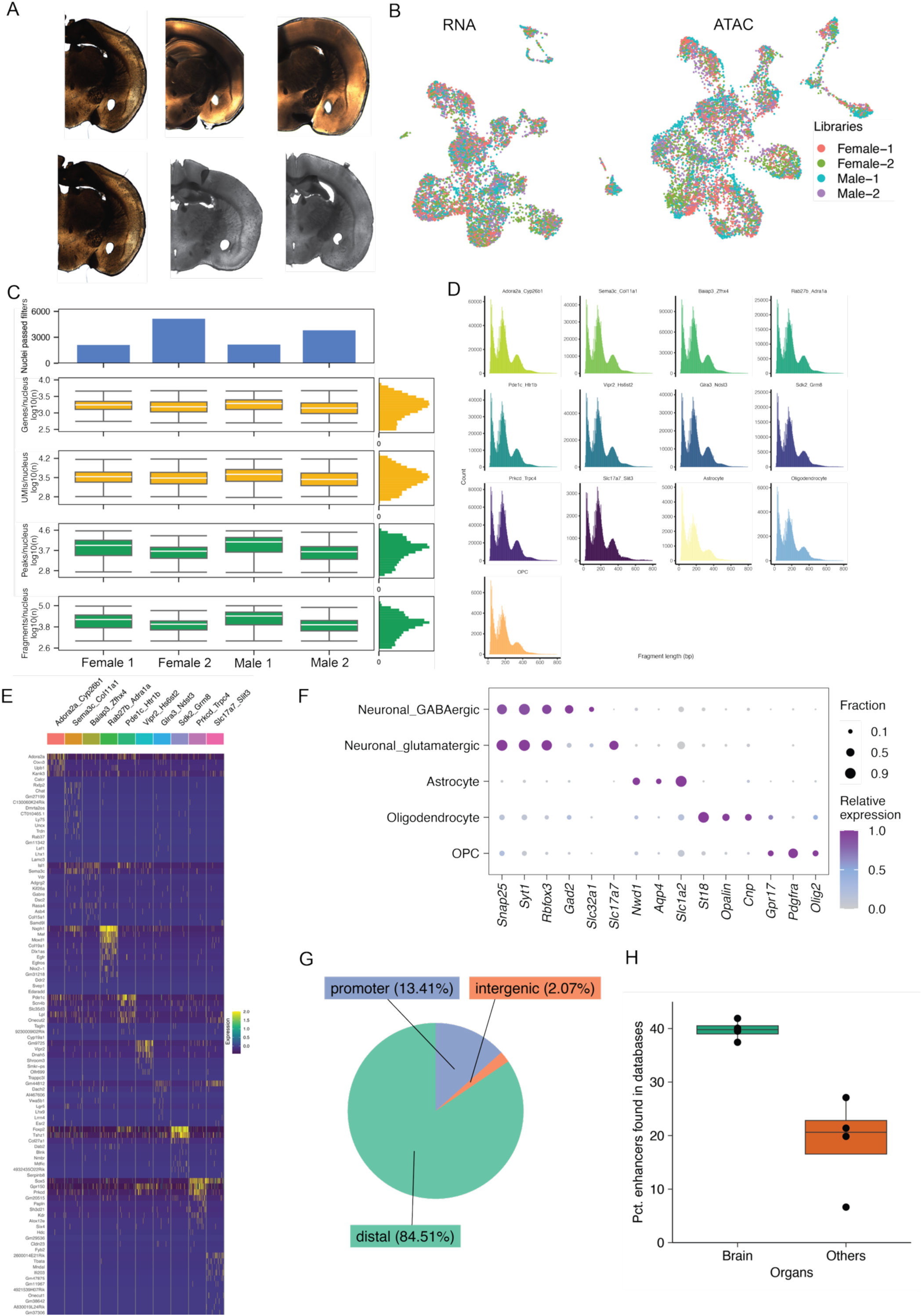
Tissue preparation and quality control for single-cell multi-omic profiling. (A) Representative brain sections showing tissue collection from amygdala regions. (B) UMAP visualization of single-cell RNA-seq (left) and ATAC-seq (right) datasets colored by sample library (Female-1, Female-2, Male-1, Male-2). (C) Quality control metrics across samples including total UMI counts, gene detection rates, and mitochondrial gene percentages. Box plots show median, quartiles, and distribution for each sample group. Histograms on right show overall distribution of quality metrics. (D) Fragment length distribution profiles for ATAC-seq libraries showing characteristic nucleosome periodicity patterns. Individual samples are color-coded, with peaks at ∼200bp intervals indicating high-quality chromatin accessibility data. (E) Comprehensive heatmap showing differential gene expression across all cell types. Rows represent individual genes, columns represent cell clusters, with hierarchical clustering revealing cell type-specific expression patterns. Color scale indicates normalized expression levels from low (blue) to high (yellow/red). (F) Dot plot showing expression levels and frequency of canonical cell type markers across all identified clusters. Circle size represents percentage of cells expressing each gene; color intensity indicates relative expression level. (G) Genomic feature annotation of ATAC-seq peaks showing distribution across promoter (13.41%), intergenic (2.07%), and distal regulatory (84.51%) regions. (H) Percent of enhancers found in our datasets represent mostly of the brain vs the other tissues.

**Figure S4. (Related to Figure 2).**
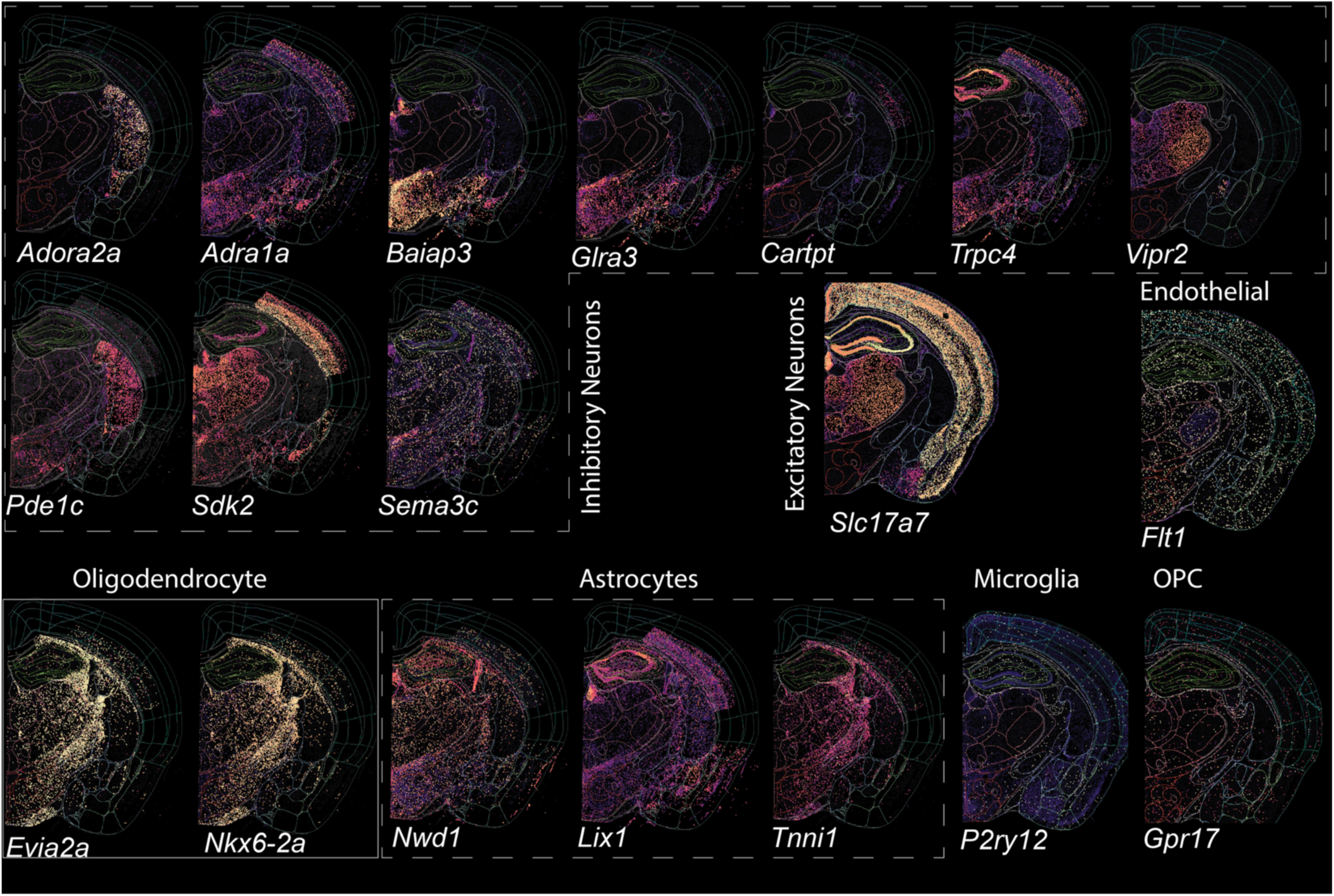
Cell type identification and marker gene expression in amygdala circuits. Spatial gene expression maps showing cell type-specific markers across coronal brain sections with identified CeA specific inhibitory, excitatory and non-neuronal cell marker genes from our snRNAseq dataset. Data source: Allen brain cell atlas (https://knowledge.brain-map.org/abcatlas).

**Figure S5. (Related to Figure 2).**
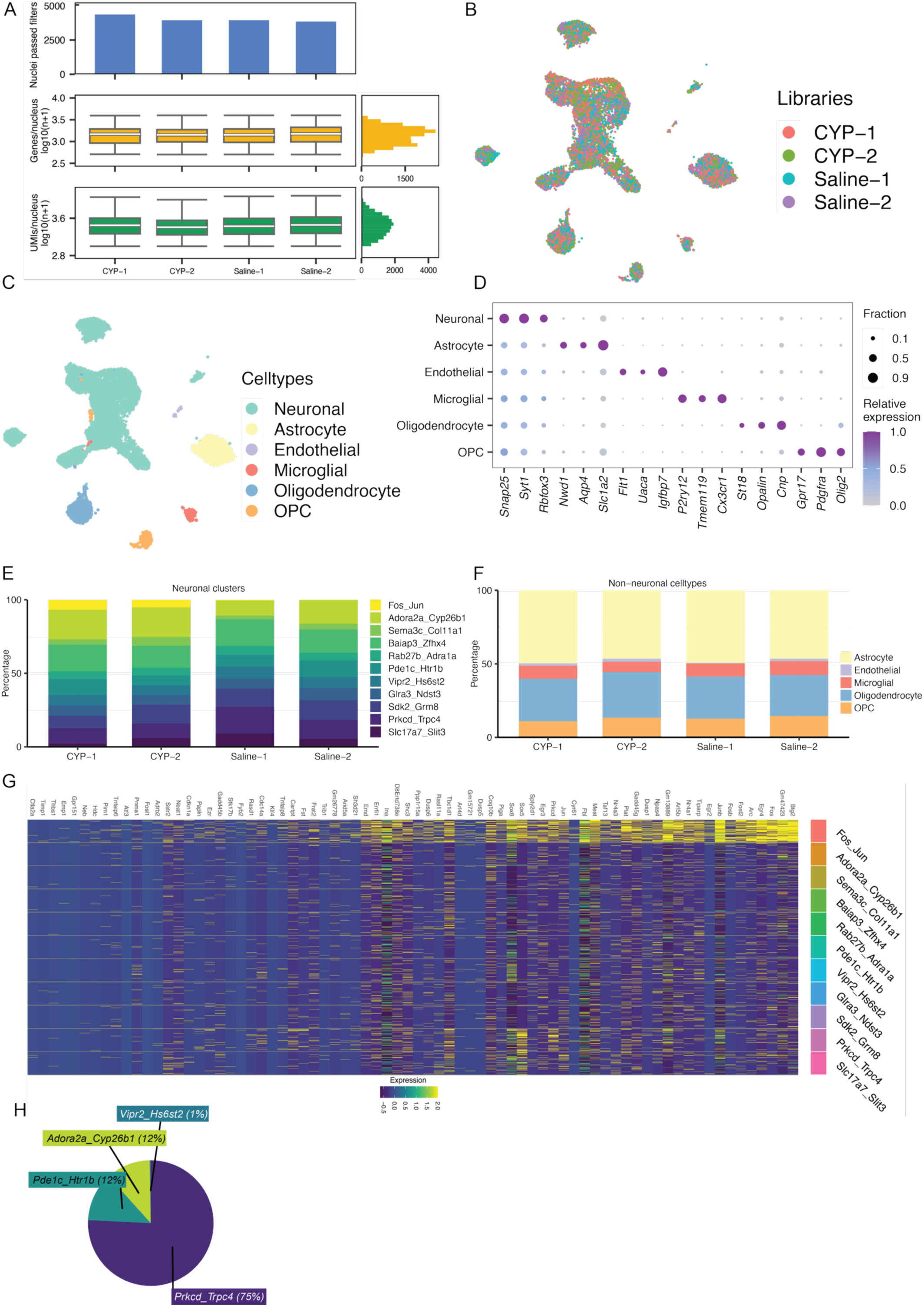
Comprehensive single-cell analysis of cell type diversity and treatment effects. (A) Quality control metrics for single-cell datasets including cell numbers, gene detection, and mitochondrial content across experimental groups. (B) UMAP visualization colored by treatment condition (CYP-1, CYP-2, Saline-1, Saline-2) showing integration of samples and cystitis-specific clustering patterns. (C) UMAP projection showing major cell type clusters identified through single-cell RNA-seq analysis, with distinct populations of neurons, astrocytes, endothelial cells, microglia, oligodendrocytes, and oligodendrocyte precursor cells (OPC). (D) Dot plot showing expression levels and frequency of canonical cell type markers across all identified clusters. Circle size represents percentage of cells expressing each gene; color intensity indicates relative expression level. (E-F) Stacked bar charts showing proportional representation of neuronal subtypes (E) and non-neuronal cell types (F) across different treatment conditions. Color coding indicates specific cell populations identified through marker gene expression. (G) Comprehensive heatmap showing differential gene expression across all cell types for saline vs CYP treatment. Rows represent individual genes, columns represent cell clusters, with hierarchical clustering revealing treatment-specific and cell type-specific expression patterns. Color scale indicates normalized expression levels from low (blue) to high (yellow/red). (H) Pie chart showing distribution of different cell-types that express Fos transcript in the CeA snRNA seq after CYP.

**Figure S6. (Related to Figure 3).**
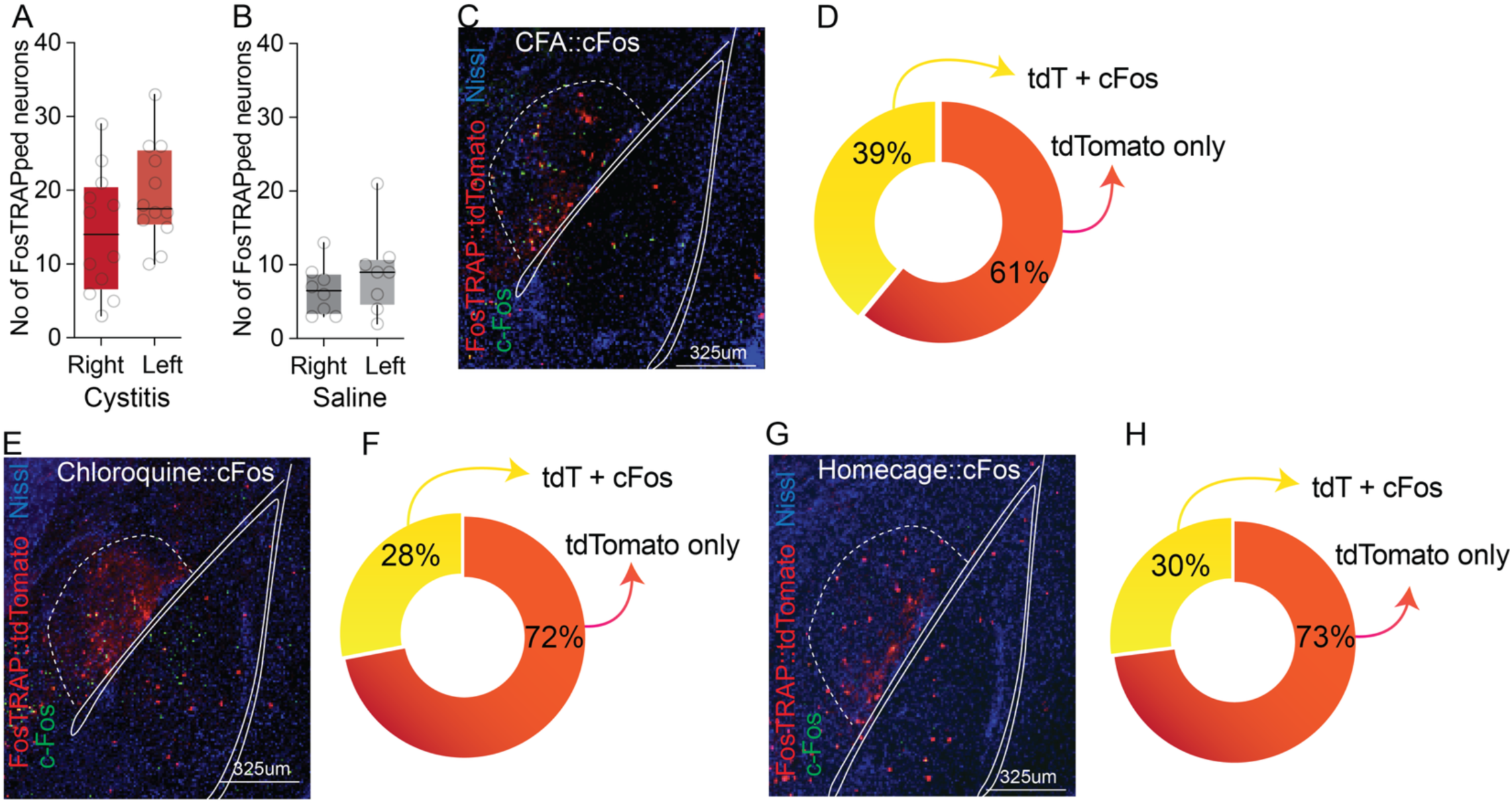
Characterization of cystitis-activated neuronal populations. (A, B) Quantification of FosTRAPed neurons in left versus right amygdala in CYP and saline TRAPed mice. n= 12/group, t-test, t=1.707, df=22, p = 0.102, n= 8/group, t-test, t=1.003, df=14, p = 0.338. (C) Representative immunofluorescence image showing colocalization of CFA-activated cFos with tdTomato^+ve^ CYP FosTRAPed CeA neurons. (D) Pie charts indicate proportion of cells relative percentages of Fos+ve neurons that are tdTomato+ve. (E) Representative immunofluorescence image showing colocalization of chloroquine-activated cFos with tdTomato^+ve^ CYP FosTRAPed CeA neurons. (F) Pie charts indicate proportion of cells overlap with chloroquine Fos+ve neurons and CYP TRAPed tdTomato+ve. (G) Representative immunofluorescence image showing colocalization of homecage-activated cFos with tdTomato^+ve^ CYP FosTRAPed CeA neurons. (H) Pie charts indicate proportion of cells overlap with homecage Fos+ve neurons and CYP TRAPed tdTomato+ve.

**Figure S7. (Related to Figure 4).**
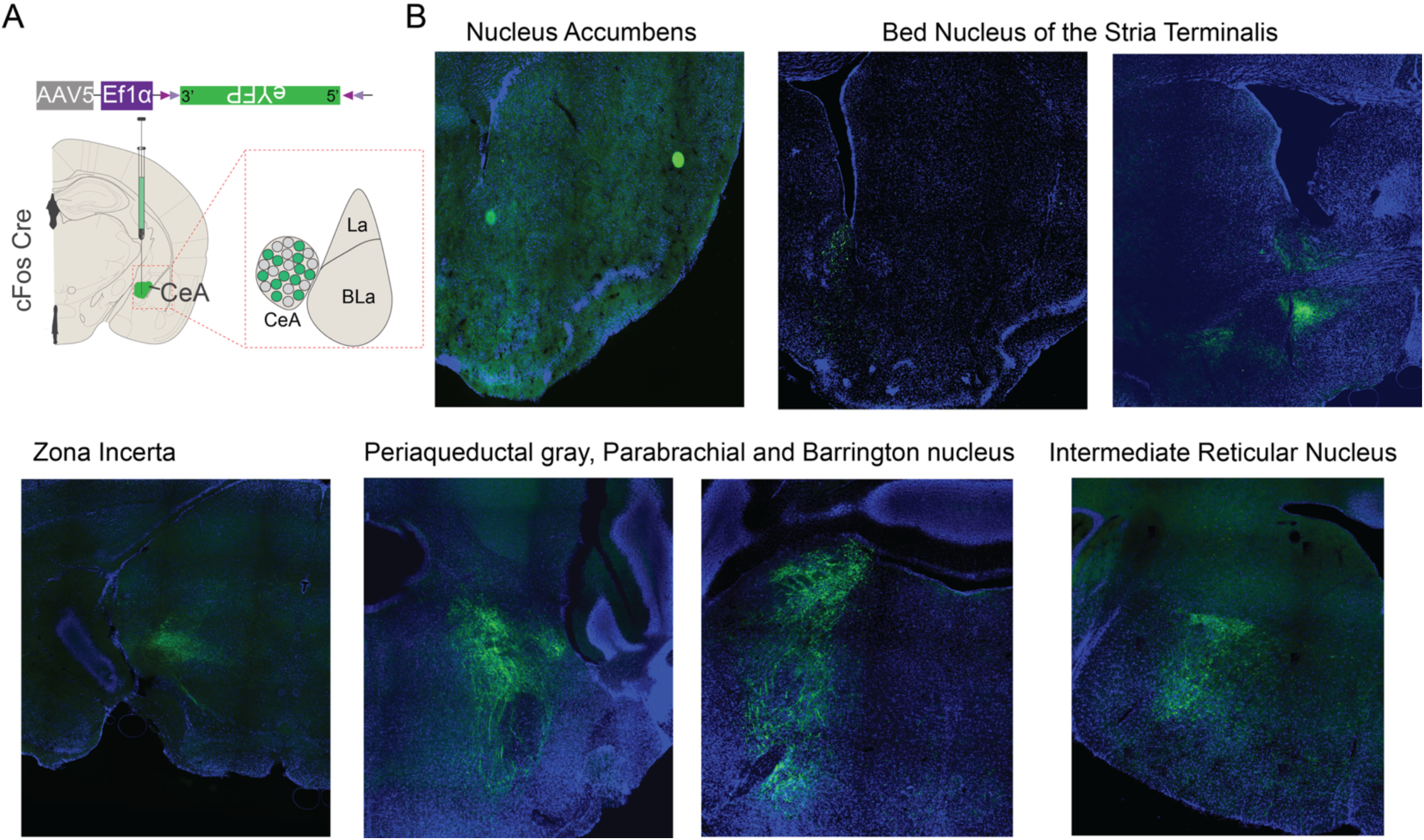
Efferent projections of the cystitis TRAPed CeA neurons. (A) Schematic of viral strategy. AAV5-Ef1α-DIO-eYFP was injected into the CeA of cFos-Cre mice to label activated neurons and their projections following stimulus exposure. Representative eYFP+ terminal labeling from CeA Fos+ neurons is observed in multiple downstream targets, including: Nucleus Accumbens, Bed Nucleus of the Stria Terminalis (BNST), Zona Incerta, Periaqueductal Gray (PAG), Parabrachial Nucleus, and Barrington’s Nucleus, Intermediate Reticular Nucleus.

